# Genomic and morphological characterization of *Trichuris incognita n. sp.*, a Human-Infecting Trichuris species

**DOI:** 10.1101/2024.06.11.598441

**Authors:** Max Alexander Bär, Nadège Akissi Kouamé, Sadikou Touré, Jean Tenena Coulibaly, Pierre Henri Hermann Schneeberger, Jennifer Keiser

## Abstract

**Background:** Trichuriasis is a neglected tropical disease that affects as many as 500 million individuals and can cause significant morbidity. For decades, it was thought to be caused by one species of whipworm, *Trichuris trichiura*. Significant differences in response rates to the best available anthelmintic treatment for trichuriasis — a combination of albendazole and ivermectin — exist across clinical trial reports yet the underlying reasons are unclear.

**Methods:** Combining long- and short- read sequencing approaches, we assembled a high-quality reference genome of *T. incognita n. sp.* isolated in an interventional study conducted in Côte d’Ivoire. The species tree of the *Trichuris* genus was constructed using 12434 orthologous groups. We sequenced a total of 747 individual worms which were used to confirm the phylogenetic placement and investigate patterns of adaptation through comparative genomic analysis.

**Findings:** Here, we present and characterize a human-infecting *Trichuris* species that we named *Trichuris incognita n. sp*. Comparative genomic analysis of genes suspected to confer resistance to either albendazole or ivermectin in helminths revealed a high number of beta-tubulin (TBB) orthologs, present in the whole population of *T. incognita n. sp.*, compared to the canonical *T. trichiura* species, but these genes were not associated to a resistant phenotype. We conducted a genome wide association study (GWAS) comparing 179 albendazole-ivermectin sensitive to 542 drug non-sensitive worms, which did not conclusively show an adaptation to drug pressure within the same species.

**Interpretation:** Our results demonstrate that trichuriasis can be caused by multiple species of whipworm and that the differences in response rates, may be a result of species responding differently to drug treatment, as opposed to an establishment of resistance. This discovery, coupled with the high tolerability of *T. incognita n. sp.* to albendazole-ivermectin marks a significant shift in how we understand and approach whipworm infections.

**Funding:** European Research Council (ERC) Nr. 101019223.

**Research in context Panel: Evidence before this study:** The recommended treatment with the benzimidazoles albendazole or mebendazole shows consistently low efficacy against *T. trichiura*, and therefore combination chemotherapy of benzimidazoles and ivermectin has been recommended. Recent clinical trials with albendazole-ivermectin, showed especially low cure rates against what was presumed to be *T. trichiura* in Côte d’Ivoire, in contrast to higher efficacy observed in Laos and Pemba, Tanzania. Amplicon sequencing data pointed towards underlying genomic differences and potentially a new species of human infecting *Trichuris* in Côte d’Ivoire.

**Added value of this study:** We identified and described a morphologically indistinguishable, novel causative agent of trichuriasis that doesn’t respond to conventional drug treatment: *Trichuris incognita n. sp*. We provide a high-quality reference genome of this novel species and sequencing data of 747 individual isolates. A genome wide association study did not conclusively show adaptation to drug pressure. This study demonstrates that trichuriasis can be caused by multiple species of whipworm and that the differences in response rates, may be a result of species responding differently to drug treatment, as opposed to an establishment of resistance.

**Implications of all the available evidence:** The current whole body of evidence calls for new effective treatments against *T. incognita n. sp*. Due to the morphological indistinguishableness, molecular diagnostic tools need to be implemented to identify species level differences as opposed to the traditional Kato Katz method. The geographical distribution of *T. incognita* as well as longitudinal changes of species distributions will need to be mapped, in addition to better understanding of the new species pathology and transmission dynamics.

## Introduction

Soil-transmitted helminthiasis is the most prevalent neglected tropical disease (NTD) worldwide. Over one billion people, particularly in impoverished nations, may develop debilitating chronic infections due to one of the five most common soil-transmitted helminths (STHs): *Ascaris lumbricoides*, *Trichuris trichiura*, the two hookworm species, *Necator americanus* and *Ancylostoma duodenale*, and *Strongyloides stercoralis*.(1) Despite their genetic diversity, the first four parasites are classified under the umbrella of STHs by the World Health Organization (WHO) and are treated with benzimidazoles, the cornerstone of large-scale deworming efforts through preventative chemotherapy (PC) since 1984 or ivermectin for strongyloidiasis.(2–5)

With more than 3.3 billion tablets of albendazole donated and distributed up until August 2020 and 5 billion treatments of ivermectin for onchocerciasis until January 2025, concern regarding anthelmintic resistance extending to humans, is rising. Reports of decreased efficacy and documented resistance to the same drugs within other helminths in livestock support projections suggesting the emergence of resistance within the next decade.(6, 7) However, no molecular evidence of resistance has yet been presented in human STHs. Notably, albendazole and ivermectin monotherapies show consistently low efficacy against *T. trichiura*, possibly due to differences in cuticle structure, mucosal embedding, or drug efflux, though these remain unproven.(8)

To circumvent potential resistance and improve efficacy against *T. trichiura*, the use of a combination therapy of albendazole and ivermectin has been taken forward. Recent clinical trials with albendazole-ivermectin, revealed especially low cure rates against what was presumed to be *T. trichiura* in Côte d’Ivoire, in contrast to higher efficacy observed in Laos and Pemba, Tanzania.(9) Amplicon sequencing data, normally not recorded in clinical trials, pointed towards underlying genomic differences and potentially a new species of human infecting *Trichuris* in Côte d’Ivoire, which may have been observed previously by S. Nissen *et al*. already in 2012.(10, 11) Despite the clinical relevance of these findings in addition to harboring information on potential immunotherapeutics,(12) genomic resources for whipworms and molecular adaptation to anthelmintic pressure in *Trichuris* remain limited.(12–14) Currently, only the genomes of three *Trichuris* species are publicly available on WormBase ParaSite,*T. suis*,(15) *T. muris*(*16*) and *T. trichiura*(16).

We present a new species of *Trichuris*, *Trichuris incognita n. sp.*, naturally infecting humans in Côte d’Ivoire. *T. incognita n. sp.* occurs where drug treatment fails and is characterized by a high number of TBB orthologs. Phylogenetic analysis based on the species tree of 12434 orthologous groups of genes, places the isolated worms within a new, monophyletic clade which is genetically closer to *T. suis* than to *T. trichiura*. We elucidate the genome of this new species, leveraging whole-genome-sequencing (WGS) data to produce a high quality reference genome of this species. Finally, we provide the first ever dataset of human infecting helminths comparing pre- and post-treatment WGS data. We investigate the whole genome sequencing data of 747 isolates in context of resistance by looking at duplication events across the species tree and conducting a GWAS.

## Methods

### Expulsion study design fieldwork

This trial was conducted in the Dabou and Jacqueville districts of Côte d’Ivoire. This area was selected, because communities with high endemicity were identified as part of a previous trial on *T. trichiura.*(9) All community members aged 6-12 who were positive for *T. trichiura* in duplicate Kato-Katz smears, with an infection intensity of at least 200 eggs per gram of stool (EPG), were eligible for trial inclusion. Excluded from the trial were individuals with reported major systemic illnesses or chronic disease, known allergy to study medications and having received anthelmintic treatment with ivermectin in the past two weeks.

This trial was conducted in accordance with the protocol, International Conference on Harmonization Good Clinical Practice E6 (R2) (ICH-GCP) and the current version of the Helsinki Declaration. Parents or guardians of participating children provided written informed consent. Children provided written assent. All authors take responsibility for the accuracy and completeness of the data and the fidelity of the trial to the protocol, which is available together with the statistical analysis plan in appendix 1. The protocol was approved by independent ethics committees in Côte d’Ivoire (reference numbers 156-22/MSHPCMU/CNESVS-kp and ECCI00918) and Switzerland (reference number AO_2022_00028).

### Treatment and sample collection from humans

Guardians/caregivers were invited to participate in an information session. The research team explained the purpose and procedures of the study, as well as potential benefits and risks of participation. The timeline of the procedures is summarized in Figure 1a. For initial diagnosis, participants were asked to provide one stool sample from which duplicate Kato-Katz thick smears using 41·7 mg of stool were prepared and assessed under a light microscope for the identification of *T. trichiura*, *A. lumbricoides*, and hookworm ova by laboratory technicians. 10% of all Kato-Katz slides were randomly chosen for subsequent quality control by picking every tenth slide of all slides read by each laboratory technician on the respective day. All eligible participants were treated with albendazole-ivermectin at day 0. Albendazole, the product of Glaxo Smith Kline (Zentel®), was a single tablet of 400 mg. Ivermectin, the product of Merck (Stromectol®), was given at a dose of 200 µg/kg. On day 8 a single oral dose of 20 mg/kg of oxantal pamoate was administered. Oxantel pamoate tablets (500 mg) were provided by the University of Basel.

**Figure 1:**
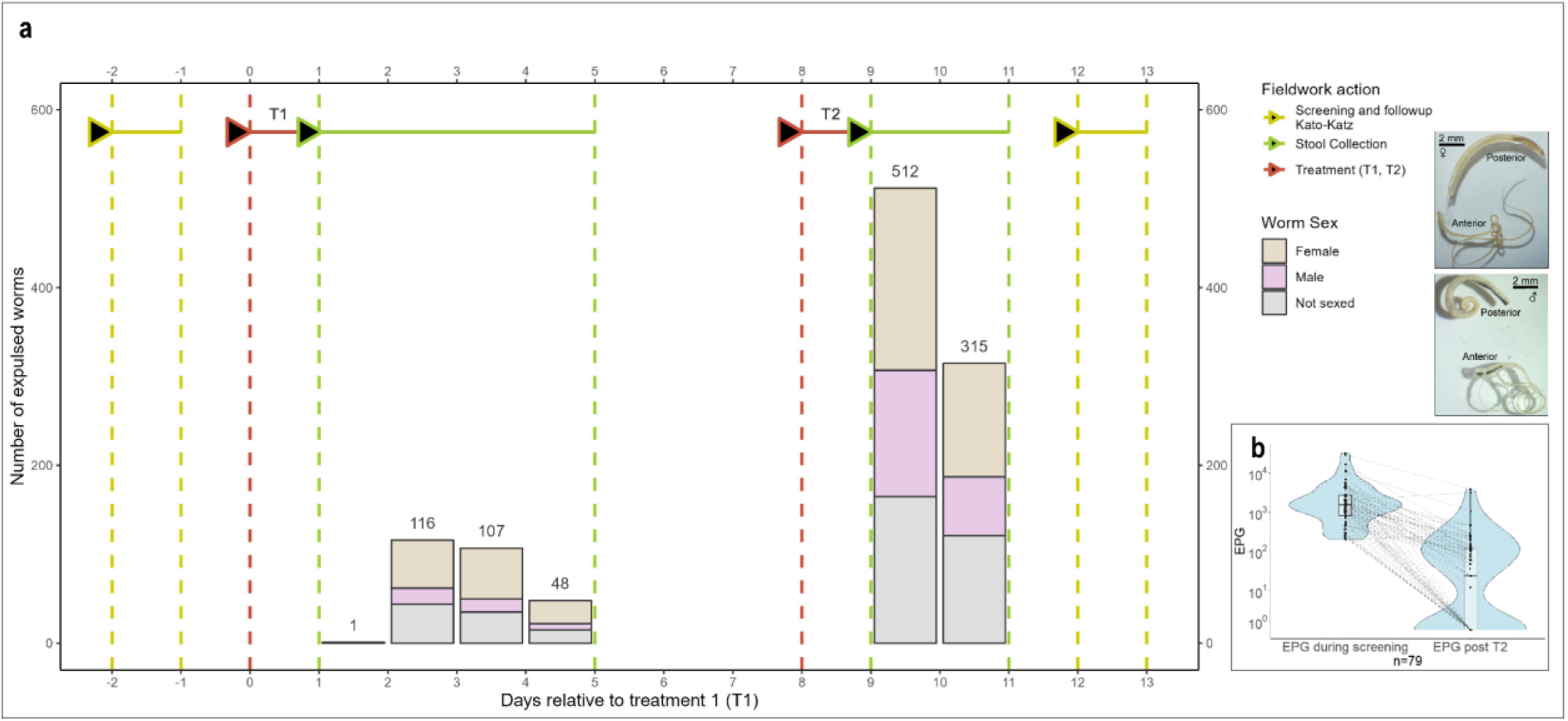
Timeline of expulsion study including the number of expelled worms and the female/male distribution of the sequenced worms a Timeline of expulsion study including the number of expelled worms and the female/male distribution. T1: albendazole-ivermectin combination treatment, standard-of-care. T2: Oxantel-pamoate treatment. Representative pictures of an expelled female and male worm are provided on the top right and bottom right respectively. Further morphological data is provided in Figure 3. b EPG before T1 and after T2. To phylogenetically place these new isolates of whole worms from humans into context of the Trichuris genus, an analysis on the mitogenome was conducted as presented in Figure 2. The phylogenetic analysis was done using the coding regions of the mitochondrial genomes of 535 individual worms that yielded a complete mitogenome and is presented in SI Figure 2. The reconstructed mitogenomes contained 13 protein coding genes (cox1-3, nad1-6, nad4L, atp6, atp8 and coB). Concatenated nucleic acid sequences from all assembled and reference mitochondrial genomes showed that the Trichuris species encountered in Côte d’Ivoire forms a separate clade from the canonical human infective T. trichiura, closer to T. suis, while the phylogenetic placement of the 16 sequenced T. suis fall within the expected clade of T. suis reference mitochondrial genomes. Previously reported sequences of a Trichuris species, isolated from a colobus monkey in Spain by Rivero et al.,{Rivero, 2020 #19} show the closest genomic relationship with the species from Côte d’Ivoire. Finally, the phylogenetic tree of the T. incognita n. sp. clade was plotted in SI Figure 3 to see if any of the clades within the species correlates to treatment, worm sex, host sex, host, coinfection with other worms, however no correlation could be identified. This species was named Trichuris incognita n. sp.

**Figure 2:**
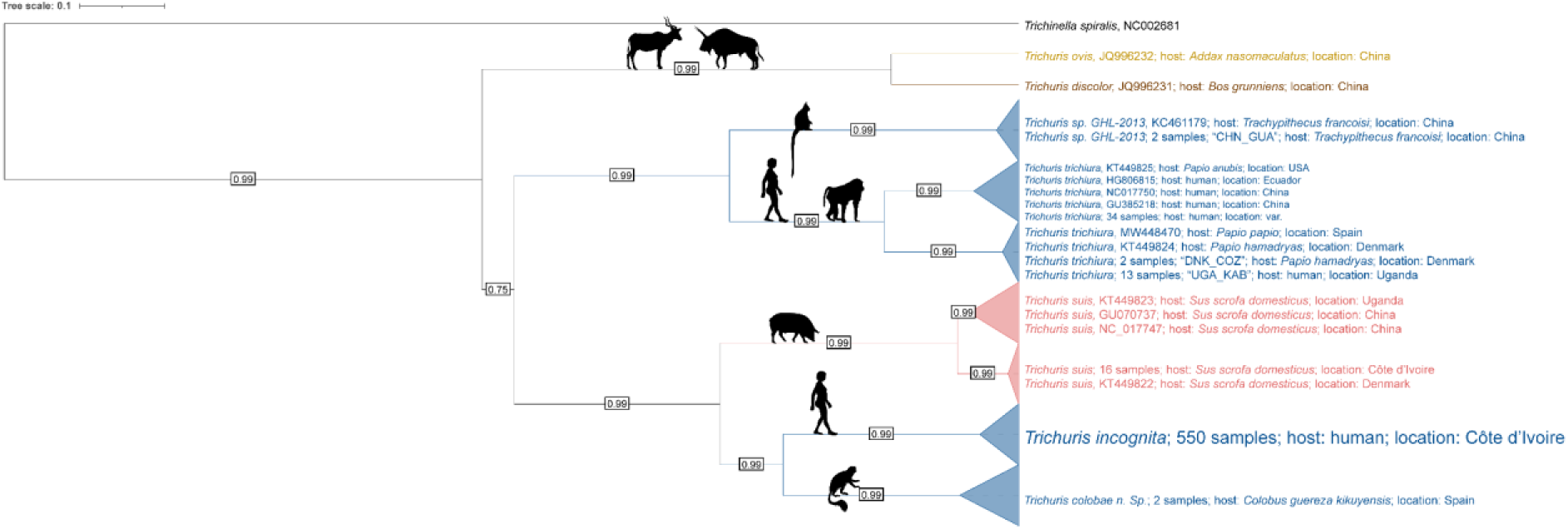
Phylogenetic placement of expelled Trichuris worms from Côte d’Ivoire

From day 1 or 2 and 8 onwards, 24h-stool was collected, based on previously reported expulsion dynamics and weighed for 2-3 days.(17, 18) Expelled worms were counted, washed twice with distilled water, once with sterile PBS and then stored in pure ethanol at 4°C. A picture of each worm in ethanol was taken and is provided on online (https://github.com/max-baer).

### Sample collection from pigs

To obtain *T. suis* worms, a list of all pig keepers was drawn by the help of local village chief. Pig keepers were gathered in formal meetings where the purpose of the study was explained. Pig keepers were asked to give written informed consent to participate in the study. Fecal samples were then collected directly from the rectum using gloves lubricated with glycerin, by a veterinarian. The samples were analyzed using the Mini Flotac technique.(19) 2 pigs that tested positive for *T. suis* roaming freely in Tiagba were sacrificed and worms were collected directly from the cecum and stored in either pure ethanol at 4 °C or mashed and stored in RNA-later at -20 °C.

### Defining a holotype for *T. incognita n. sp.* and depositing a sample at the natural history museum in Basel

A female specimen of *T. incognita n. sp.* isolated from *Homo sapiens* in the town of Tiagba in the district des lagunes in Côte d’Ivoire has been chosen as the holotype and was archived at the natural history museum in Basel under the number NMB-79 a in absolute ethanol together with a male paratype. All authors of this publication have been registered as the discoverers of this new species. This work and the nomenclatural acts it contains for the new species have been registered in ZooBank: LSID: 34409100-BF94-4A0D-9088-D28215D231F6.

### DNA extraction and high-throughput sequencing

Worms stored in absolute ethanol were photographed and used for sequencing experiments. The posterior end of the worms was severed and DNA was extracted using a DNeasy Blood & Tissue Kit (Qiagen, Cat: 69504) according to the manufacturer’s protocol from the anterior part of the worm. Sequencing libraries for illumina were prepared using the NEBNext® Ultra™ II FS DNA Library Prep Kit (New England Biolabs, Cat: E7805L) according to the manufacturer’s protocol after which DNA concentration and library size was determined (HS NGS Fragment Analyzer). Libraries were sequenced using 150 bp paired-end chemistry at an average 20X coverage on an Illumina NovaSeq6000 platform with an S4 flow cell at the Genomics Facility Basel as part of the Department of Biosystems Science and Engineering. Sequencing libraries for Nanopore were prepared using the Native Barcoding Kit 24 V14 (Oxford Nanopore Technologies, Cat: SQK-NBD114.2) from a single whole female worm and sequenced on a PromethION flow cell and PromethION 2 device.

### PCR and Nanopore sequencing experiment of the TBB gene

Two primers were selected surrounding the regions containing known mutations (Phe168, Glu198, Phe200), observed variable position (Ser194) and predicted truncation site. The primer sequences were AAAGAGACCGGACATTTCGC for the forward primer and TGAATTGCCTGGTTTCTAGAATG for the reverse primer and the amplified region was 402 base pairs. Genomic DNA from one sequenced worm was used as input and the OneTaq® One-Step RT-PCR Kit was used to amplify the region during 40 cycles with an annealing temperature of 56°C for 30 seconds and an extension temperature of 68°C for 1 minute Sequencing libraries for Nanopore were prepared using the native barcoding kit 24 V14 (Oxford Nanopore Technologies, Cat: SQK-NBD114.2), and sequenced on a Flongle flow cell and the MinION Mk1c device. The dorado basecaller (version 0.7.2, model 400bps_hac@v5.0.0) was used for duplex basecalling. Reads were filtered by alignment score (AS>600) and length (>390 BP) and aligned using minimap2 to the reference TBB sequence. Variants were called using bcftools and coverage plots were created in R.

### Pipeline for raw read processing and mitochondrial genome assembly

For all sequencing, assembly and annotation pipelines Nextflow (version 23.04.1 build 5866) was used with Java (version 11.0.3).(20). Raw reads were first processed using Trimmomatic (version 0.39).(21) for PE reads, adapters were removed and N-bases trimmed. Next, reads were channeled to getorganelle (version 1.7.7.0) to reconstruct the mitochondrial genome.(22) Runs which resulted in a complete mitochondrial genome were further processed by flipping in cases where the reverse complement was generated and adjustment to COX1 as starting gene. Reference and newly constructed mitochondrial genomes were annotated and visualized using MitoZ (version 3.6).(23) Coding genes were then extracted and aligned as amino acid and nucleic acid sequences using MAFFT (version 7.490).(24) All reconstructed mitochondrial genomes and the aligned MAFFT files used as input for phylogenetic inference are provided online.

### Phylogenetic analysis

BEAST2 (version 2.7.5) was used for all phylogenetic inference with a pure birth process (Yule process).(25) Aligned concatenated nucleic acid sequences from all assembled and reference mitochondrial genomes were treated as homochronous. The JC69 substitution model with gamma-distributed rate heterogeneity was used (JC69 + Γ4) and a strict molecular clock was assumed. Trees and clock models were linked and all model parameters were estimated jointly. A Markov Chain Monte Carlo was run for each analysis. Tracer (version 1.7.2) was used to assess convergence and effective sample size. The percentage of samples discarded as burn-in was at least 10%. For the amino acid inferred species tree, 2 assembled sequences of each subclade clade with a posterior distribution over 0.95 were selected at random. The mito REV substitution model was used and all genes were partitioned separately, allowing for different rates for each gene. Effective sample size (ESS) was at least 900 for each of the inferred parameters. The intraspecies phylogeny was inferred using a TN96 + Γ4 model. Each gene was partitioned separately and codon positions 1 and 2 were partitioned separately to 3. The mean mutation rates of the genes and there codon positions are provided in the supporting information and show a faster mutation rate for codon position 3 in all cases.

### Pipeline for Hybrid-genome assembly and gene prediction

Nextflow (version 23.04.1 build 5866) was used with Java (version 11.0.3). Nanopore raw reads from a single worm were channeled into chopper (version 0.6.0) where reads below 5kb and low quality reads were filtered out and adapters trimmed.(26, 27) Trimmed reads were channeled into Kraken2 (version 2.1.1) and reads with a contamination ratio >0.5 were filtered out with a custom R (version 4.3.0) script, resulting in a 50X coverage.(28) Illumina reads of the same worm at a 65X coverage were channeled into Kraken2 where reads of impurities were identified and filtered out. Illumina and Nanopore channels were combined to run a hybrid de-novo genome assembly using MaSuRCA (version 4.0.9) which allowed for paired end illumina reads and nanopore reads.(29) A total of 1564 contigs were assembled and the raw assembly is provided on github. The BUSCO showed a completeness of 64.3% of complete BUSCOs, 11.2% fragmented and 24.5% missing and the N50 was 0.26 Mb. The outputted contigs were channeled into RagTag (version 2.1.0) for correction and scaffolding using the *T. trichiura* (PRJEB535) reference genome.(30) The corrected scaffolds were then assigned to one of three linkage groups from *T. muris* using Liftoff (version 1.6.3),(31) as described in literature.(32) The genome was frozen for quality assessment using BUSCO (version 5.1.2 with Metazoa_obd10 database)(33) and QUAST (version 5.0.2).(34) At this point all available reference whole genomes of *Trichuris* species and *Trichinella spiralis* were channeled into the pipeline to run a de-novo gene prediction on all reference genomes and reduce methodological bias from gene prediction software. Repeat regions were masked using RepeatMasker (version 4.1.4) before channeling the genome to BRAKER (version 3.0.6, Braker2 pipeline using the Metazoa protein database from OrthoDB 11).(35, 36) Gene prediction statistics including exon and intron count, mean length and median length were gathered using a custom bash script.

### Phylogenetic orthology inference, species tree, orthologous group functional annotation and GO term enrichment analysis

Orthofinder (version 2.5.5) was used to infer orthology and the species tree of the braker2 predicted genes.(37) 81425 out of 92202 (88.3%) genes were classified into 12434 orthologous groups with a mean group size of 6.5 and median group size of 5. Gene ontology of orthologous groups of interest was inferred using InterProScan (version 5.63-95.0). A list of resistance associated genes was downloaded from WormBase and blasted against all orthologous groups using an E-value threshold of 1e^-50^ to identify orthologous groups of resistance associated genes. In SI Table 4 a list of all genes with matching orthologous groups and Gene IDs from WormBase ParaSite is provided. Species tree with heat map was visualized using iTOL (version 6.8.1).(38) Duplicated, gained, lost and retained genes across the species tree were inferred using OMA standalone (version 2.6.0) and pyHam (version 1.1.11).(39, 40) Massively expanded gene families were defined as ≥7 copies in *Trichuris incognita n. sp.* and ≤4 in all other species and a list is provided in the supporting information together with the functional annotation. The orthology of predicted transcripts was inferred analogously. A custom R-script was used to visualize the Venn diagram and identify transcripts shared by *T. incognita n. sp.*, *T. trichiura* and *T. muris*. SI Table 3, a list of these transcripts with their functional annotation is provided. Protein structures were predicted using Alphafold (version 2.2.0, template date cutoff was set to 2021-09-01) and visualized with PyMOL (version 2.5). GO terms were extracted from the InterProScan annotated genes file of *T. incognita n. sp.* (background GO terms) and of *T. incognita n. sp.* genes shared with *T. trichiura* but not with *T. suis* or *T. muris* (GO terms of interest). A fishers exact test was performed on the GO term counts of each GO term in the intersection compared to the background GO terms. All gene lists of *T. incognita n. sp.* and the InterProScan annotations are provided on github.

### Pipeline for variant calling

Nextflow (version 23.04.1 build 5866) was used with Java (version 11.0.3). Variants were called by adapting the analytical pipeline in by Doyle *et al.* into Nextflow .(32) Raw reads were first processed using Trimmomatic (version 0.39).(21) BWA (version 0.7.17) and SAMtools (version 1.14) were used to index the assembled reference genome, and align, sort and filter raw reads.(41) Flagstat and Kraken2 (version 2.1.1) were used as QC tools. The coverage ratio was used to classify the gender of each worm by analyzing the coverage ratio of the largest X-chromosome scaffold to the largest scaffold on chromosome 2. Next, the standard GATK (version 4.2.6.1) pipeline was followed for variant calling, base quality score recalibration (BQSR) and genotyping. First, duplicates were marked using the MarkDuplicatesSpark option in GATK. Basecalling was done in a first round to obtain a subset of very likely SNPs for base quality score recalibration. Plots were generated for QUAL, DP, QD, FS, MQ, MQRankSum, SQR, ReadPosRankSum. Heterozygosity was set to 0.015 and indel heterozygosity to 0.01.(32) Initial hard filtering of the variants for BQSR was done in a way to exclude 5% of SNVs with the following lowest quality metrics: QUAL < 194 || QD < 10.3 || MQ < 43.61 || FS > 9.9 || SOR > 2.30 || MQRankSum < -1.269 || ReadPosRankSum < - 1.174 for SNPs and QUAL < 101 || QD < 5.3 || FS > 11.4 || ReadPosRankSum < -1.336 for indels. Next, BQSR was conducted before generating a merged gVCF which was used for genotyping and then filtered for SNVs using the following hard filter expression: QUAL < 35 || QD < 1.3 || MQ < 28.76 || FS > 50.5 || SOR > 2.30 || MQRankSum < -4.261 || ReadPosRankSum < -1.421 for SNPs and QUAL < 32 || QD < 0.5 || FS > 30.1 || ReadPosRankSum < -1.836 for indels. Indels were filtered out and the variants were further filtered using the genotype filter expression DP < 3 and a minor allele frequency of 0.02 which resulted in a total of 6’282’017 SNPs. Next, genes in the VCF file were annotated using BCFtools (version 1.15) and BEDTools (version 2.30.0) and the braker2 produced gff file containing all genes of *T. incognita n. sp.*. Using SnpEff (version 5.2c) a total of 111’994 missense SNPs leading to a different amino acid on the genes of *T. incognita n. sp.* . This set of SNPs as well as the non-filtered set was used as an input in the GWAS.

### Pipeline for genome wide association study

Plink (version 1.9) was used to conduct the genome wide association study.(42) First, the principal components were analyzed, as presented in SI Figureto assess population stratification and correct any falsely assigned sexes. Next, the phenotype data (SI Table 8) on treatment one (T1) or treatment 2 (T2) was added using the “--pheno” option as well as the worm sex using the “--update-sex” option. Next, the complete set of variants, as well as the subset of variants leading to missense mutations on predicted genes were filtered using a minor allele frequency of 0.05, a missing genotype frequency of 0.05, and a deviation from the Hardy-Weinberg Equilibrium at a significance level of p=0.001 resulting in 1254247 variants over all, and 56273 missense variants on predicted genes, which were used for association testing. The GWAS was performed a total of 24 times and the results of each run are presented to be as unbiased as possible. Each GWAS for both variant sets was performed once for both sexes together, and once for each sex individually, to account for variation associated to worm sex. Next, each GWAS was performed once with population stratification and once without population stratification as the PC1 and PC2 did not show any significant association to treatment, however, did show association to some villages in both male and female populations. Finally, the GWAS was ran once with a logistical model and once with a linear model resulting in 24 runs of the GWAS. The GWAS’ were ran using linear and logistic models with the “--assoc" or “--logistic” option respectively, with both sexes, only females and only males. Next, a principal component analysis was conducted to account for population stratification. The option “--pca" was used on the file containing all filtered variants. As an example for the subset of variants on coding genes, which is representative for all runs, the analysis of the principal components revealed an eigenvalue of 52.7 for the first principal component and 3.1 for the second, indicating that the consideration of only the first principal component will account for most population stratification. Next, the logistic and linear models were run again under consideration of the first principal component by using the options “-- covar" and “--covar-number 1” and the “--linear" or “--logistic” option for the linear or logistic model respectively for both sexes, only females and only males respectively on both sets of variants. To visualize the results and generate the QQ-plots the R package qqman (version 0.1.9) was used.

## Data availability

Raw sequencing data of 747 samples together with the associated metadata and the whole genome of *T. incognita* (TaxID: 3388467) is available under the Sequencing Read Archive, BioProject “PRJNA1282940”. 551 assembled mitochondrial genomes including *T. incognita n. sp.* and *T. suis*, as well as reference mitochondrial genomes and the *de novo* assembled *Trichuris incognita n. sp.* reference genome are available under the BioProject number and on GitHub https://github.com/max-baer. The publicly available mitochondrial genomes used for comparative analysis are available from ENA and NCBI. There are no restrictions on data availability.

## Code availability

Custom code to analyze data and reproduce the figures presented is available at https://github.com/max-baer. Where available, all input scripts and log files are provided including nextflow log files documenting a complete run, BEAST2 XML input and log files.

## Results

### *Trichuris* expulsion study confirms a new species of whipworm infecting humans, responding poorly to conventional albendazole-ivermectin treatment

We screened 670 children in 7 villages, namely Akakro, Ahouya, N’Doumikro, Tiagba, Bekpou and Teffredji in southern Côte d’Ivoire for trichuriasis. 243 participants were enrolled in the study and received a single dose albendazole-ivermectin combination treatment (T1) after which stool was collected daily and screened for worms for up to four days. Subsequently, 8 days after T1, the same individuals received a single oral dose of the trichuricidal drug oxantel- pamoate (T2) after which stool was collected for 2 consecutive days, as presented in Figure 1a. We isolated 271 worms after T1, indicating partial efficacy of the combination treatment, with an expulsion peak on days 2 and 3 and 827 worms after T2 with expulsion peaking on day 1 and 2 after treatment. A total of 1098 *Trichuris* worms were collected, of which 747 provided enough DNA for genomic analyses. The female to male ratio of the worm sex was 3.42:1 and 1.6:1 during T1 and T2 respectively. A significant correlation (R^2^ = 0.34; *P* = 0.0061, SI Figure 1) was observed between the number of expelled worms and the mean egg count per day, yielding a fecundity of 14307 eggs per female worm per day, as presented in SI Figure 1. Figure 1b shows near complete expulsion of all worms as the median EPG dropped from 1524 to 24 at the follow-up time point on day 12. To assess the genetic relatedness of human worms with *T. suis*, 25 individual worms from 2 swine hosts were collected directly from the caecum. A picture of each worm which was used to identify the worm sex is provided on github.

Bayesian inferred phylogenetic tree based on the amino acid sequences of genes in the mitochondrial genomes of 535 *Trichuris incognita n. sp.* worms and available reference mitochondrial sequences, using *Trichinella spiralis* as an outgroup. Timescale indicates numbers of substitutions per site.

Morphological description of *T. incognita n. sp*.

This parasite has a threadlike structure, filiform. Both female and male can be divided into a thin hair-like anterior part and broader handle-like posterior section. The posterior section of the female worms is slightly ventrally incurved compared to the strong ventral incurvature of the male, adopting a Fibonacci spiral-like structure. The overall length of the worms (displayed in Figure 1a) is 45 mm and 49 mm and the ratio of anterior to posterior is 2.5 and 3.1 for female and male worms respectively. An analysis of a subset of five male and five female worms are within literature reported min-max values for *T. trichiura* lengths as shown in SI Table 1.(43) The main organs of the male posterior end are the intestine, testis and ejaculatory duct, which run in parallel along the long axis of the body (Figure 3a). At the height of the ending of the testis, the ejaculatory duct and intestine join, forming the cloaca, which opens at the posterior end of the male body. The distal cloacal tube contains the spicule. The spicule is surrounded by a shiny spicule sheath (Figure 3b, 3c) with a granular surface structure. The female posterior starts at the esophagus-intestinal junction, where the vulva is located (Figure 3d). The vagina shows thick walls, which connect back into the oviduct (Figure 3e). The cuticle of the anterior structure shows two patterns, one, which is striated with transverse grooves on one side and the other, a tuberculate band (Figure 3g). Within the anterior part is the stichosome, comprised of a row of stichocytes surrounding the esophagus which starts at the mouth (Figure 3i), and continues to the posterior part of the worm merging into the intestine (Figure 3d) and marking the division between anterior and posterior section. The ova present in the oviduct contained two polar plugs, were 68 μM long and 29 μM wide (Figure 3f) placing them closer to a larger observed egg population of two distinct ova groups of *T. trichiura* in literature.(44)

**Figure 3:**
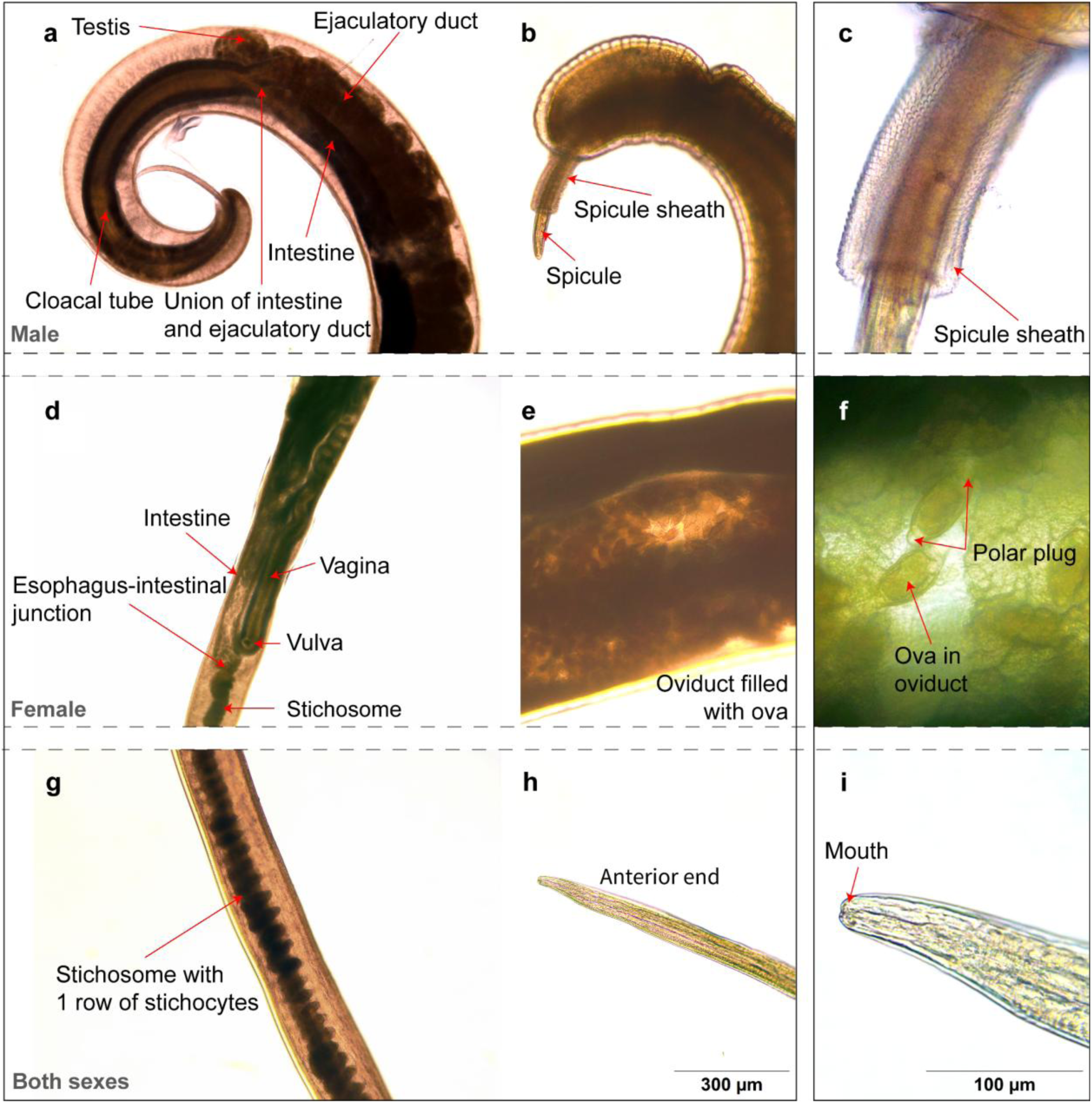
Morphology of T. incognita n. sp. the same scale was used for a,b,d,e,g and h, and a different one for c, f and i and is provided in the bottom right corner. a Posterior end showing male organs including testis, ejaculatory duct, intestine, union of intestine and ejaculatory duct, cloacal tube and spicule with a ruptured sheath. b Male spicule with an intact sheath partially covering the spicule. c 40X magnification of the shiny spicular sheath showing a surface structure. d Morphology showing female reproductive organs at the intersection between anterior and posterior part of the worm including stichosome, esophagus-intestinal junction, intestine, vagina and vulva. e Enlarged oviduct filled with T. incognita n. sp. ova. f 40X magnification on ova showing polar plugs. The ova are 68 μM long and 29 μM wide in this image. g Anterior showing one row of stichocytes and 2 types of cuticle patterns. The left side showing transverse grooves and the right side the tuberculate band. h Anterior ending leading up to the mouth. i 40X magnification of the mouth.

**Table 1:**
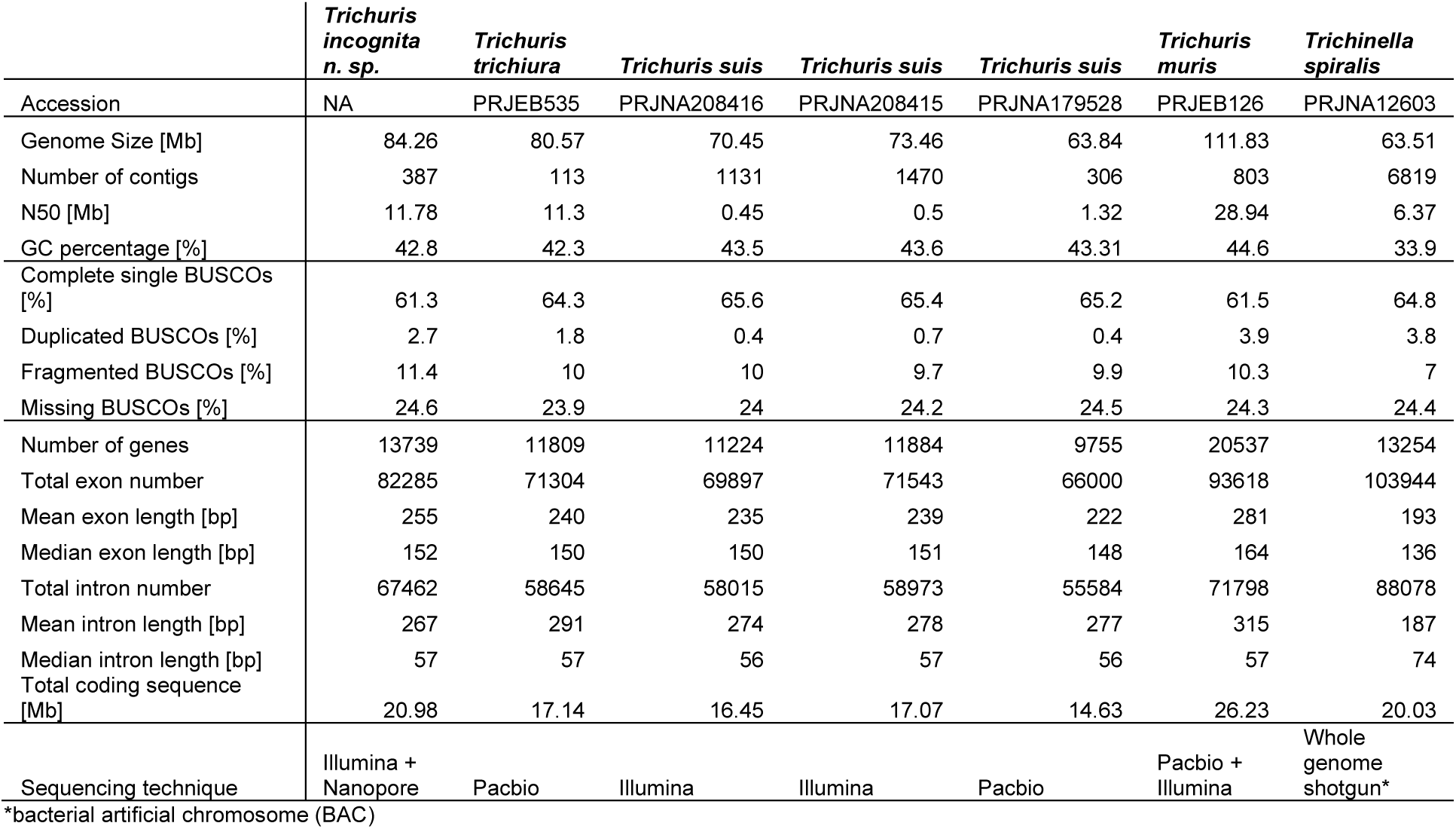
Whole genome assembly metrics.

|  | *Trichuris incognita n. sp.* | *Trichuris trichiura* | *Trichuris suis* | *Trichuris suis* | *Trichuris suis* | *Trichuris muris* | *Trichinella spiralis* |
| --- | --- | --- | --- | --- | --- | --- | --- |
| Accession | NA | PRJEB535 | PRJNA208416 | PRJNA208415 | PRJNA179528 | PRJEB126 | PRJNA12603 |
| Genome Size [Mb] | 84.26 | 80.57 | 70.45 | 73.46 | 63.84 | 111.83 | 63.51 |
| Number of contigs | 387 | 113 | 1131 | 1470 | 306 | 803 | 6819 |
| N50 [Mb] | 11.78 | 11.3 | 0.45 | 0.5 | 1.32 | 28.94 | 6.37 |
| GC percentage [%] | 42.8 | 42.3 | 43.5 | 43.6 | 43.31 | 44.6 | 33.9 |
| Complete single BUSCOs [%] | 61.3 | 64.3 | 65.6 | 65.4 | 65.2 | 61.5 | 64.8 |
| Duplicated BUSCOs [%] | 2.7 | 1.8 | 0.4 | 0.7 | 0.4 | 3.9 | 3.8 |
| Fragmented BUSCOs [%] | 11.4 | 10 | 10 | 9.7 | 9.9 | 10.3 | 7 |
| Missing BUSCOs [%] | 24.6 | 23.9 | 24 | 24.2 | 24.5 | 24.3 | 24.4 |
| Number of genes | 13739 | 11809 | 11224 | 11884 | 9755 | 20537 | 13254 |
| Total exon number | 82285 | 71304 | 69897 | 71543 | 66000 | 93618 | 103944 |
| Mean exon length [bp] | 255 | 240 | 235 | 239 | 222 | 281 | 193 |
| Median exon length [bp] | 152 | 150 | 150 | 151 | 148 | 164 | 136 |
| Total intron number | 67462 | 58645 | 58015 | 58973 | 55584 | 71798 | 88078 |
| Mean intron length [bp] | 267 | 291 | 274 | 278 | 277 | 315 | 187 |
| Median intron length [bp] | 57 | 57 | 56 | 57 | 56 | 57 | 74 |
| Total coding sequence [Mb] | 20.98 | 17.14 | 16.45 | 17.07 | 14.63 | 26.23 | 20.03 |
| Sequencing technique | Illumina + Nanopore | Pacbio | Illumina | Illumina | Pacbio | Pacbio + Illumina | Whole genome shotgun* |

\*bacterial artificial chromosome (BAC)

### *T. incognita n. sp.* hybrid *de novo* whole genome assembly and comparative genomic analyses

The *de novo* assembled genome of *T. incognita n. sp.,* sequenced at a coverage of 50X Nanopore and 65X Illumina reads, resulted in a length of 84.26 Mb, compared to 80.57 Mb of *T. trichiura*, 63.84 Mb, 73.46 Mb and 70.45 Mb of *T. suis* and 111.83 Mb of *T. muris*. The *T. incognita n. sp.* genome consisting of 387 contigs with an N50 of 11.78 Mb is presented in Table 1. The genome completeness amounted to 75.4% of whole, duplicated, and fragmented single copy orthologues as determined by BUSCO. GC content was 43%. Genes were predicted *de novo* for all available reference whole genomes using the braker2 pipeline and the metazoa database, resulting in the prediction of 13739 genes for *T. incognita n. sp*. An exploratory analysis of these genes in context to the *in silico* predicted genes of *T. trichiura*, *T. suis* and *T. muris* presented in SI Table 2 and SI Table 5 identified one family of each serpins (OG0000305) and chymotrypsin-like serine proteases (OG0010319) that are shared only by *T. trichiura* and *T. incognita n. sp.* (SI Figure 4) additionally to a family of tetraspanins (OG0009162) also shared by *T. muris*.(45, 46)

**Figure 4:**
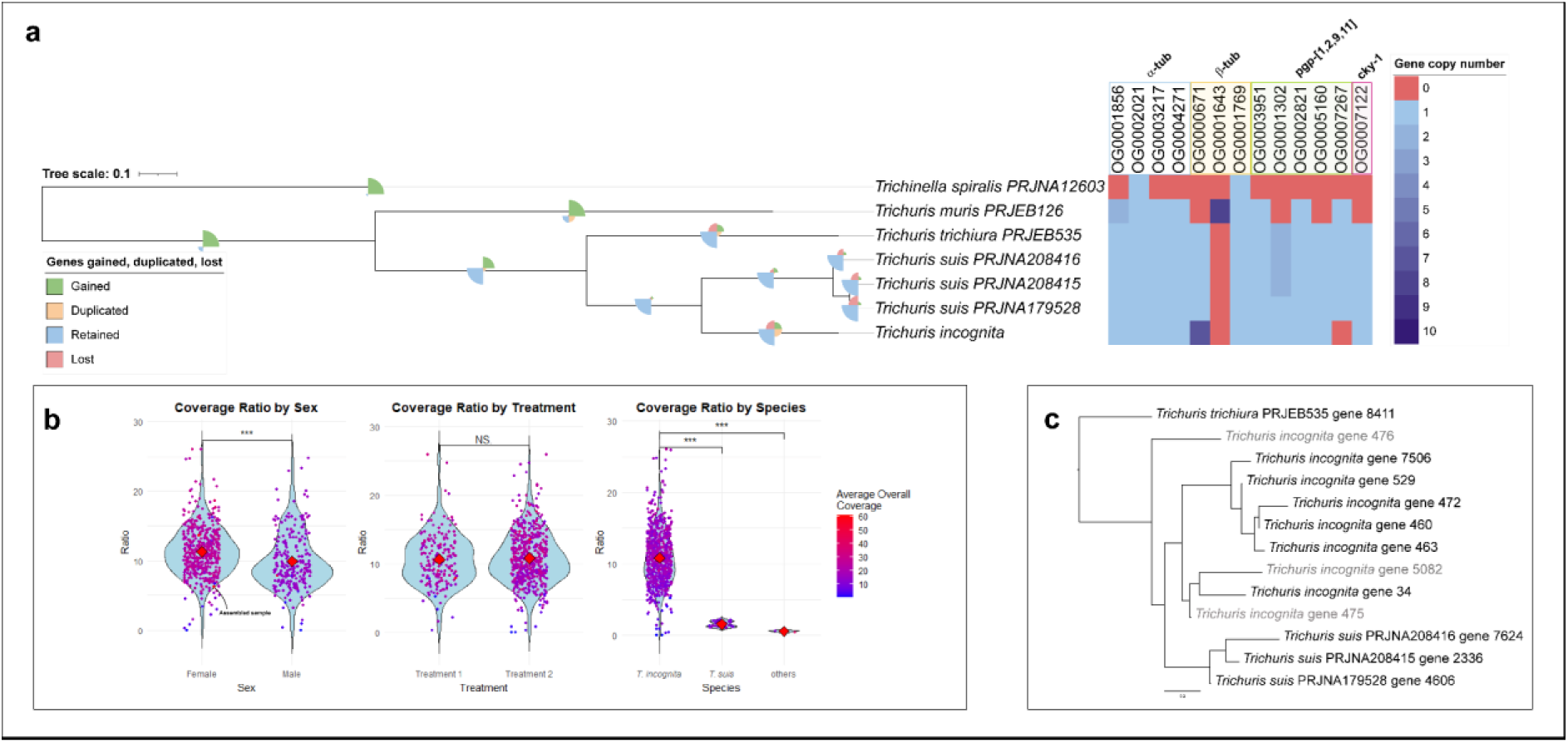
Duplications across the species tree show a high number of β-tubulin gene orthologs. a Species tree indicating gene duplication, gains, losses and retention as pie charts across the tree. Heat map of ortholog numbers for all genes present in orthologues groups of genes, which have previously been associated to resistance and the α-tubulin gene. b Coverage ratio compared between worm sex (n=747), treatment (n=747) and from T. incognita n. sp. (n=747) to T. suis (n=26) from Côte d’Ivoire and other species. Three T. suis genomes and one T. trichiura genome were used to calculate the average coverage in the species containing only one gene in OG0000671. *** signifies a p-value < 0.001 from a two tailed t-test. c TBB gene tree of OG0000671, timescale indicates numbers of substitutions per site.

### The species tree confirms the existence of a new species of *Trichuris*, and reveals highly duplicated genes associated to microtubule dynamics and mitotic processes

The species tree of the *Trichuris* genus presented in Figure 4a was inferred using species tree inference from all genes (STAG)(47, 48) with 12434 orthologous groups of the braker2 predicted genes in the publicly available whole genomes of *Trichuris* species, *T. incognita n. sp.* and *Trichinella spiralis*. The species tree shows the same topology as for the mitogenome inferred phylogeny.

We further inferred the gene orthology of all genes predicted in the publicly available whole genomes of *Trichuris* species, *T. incognita n. sp.* and *Trichinella spiralis* to identify duplication events of genes that have been associated with resistance to either albendazole or ivermectin in helminths. These resistance-associated genes include TBB for albendazole or glutamate-gated chloride channel subunits (*glc*-1, *glc*-2, *glc*-3, *avr*-14, *avr*-15), voltage gated chloride ion channels (*clh*-3), sodium:potassium:chloride symporter (*nkcc*-1), bestrophin chloride channel (*best*-19), p-glycoprotein ABC transporters (*pgp*-1, *pgp*-2, *pgp*-9, *pgp*-11), and transcription factor *cky*-1, for ivermectin respectively.(14, 49) Amongst these genes, one orthologuous group (OG0000671) of the TBB gene showed 9 orthologs of beta tubulin gene in *T. incognita n. sp.* and only 1 ortholog in all *T. suis* and *T. trichiura* reference species as presented in Figure 4a. One gene was fragmented into 2 sections, g475 and g476, and g5082 was also found to be fragmented. Intriguingly, the α-tubulin gene was only present as a single ortholog. SI Figure 5 shows the 7 predicted TBB genes from the braker 2 pipeline. None of the genes show previously encountered Phe168, Glu198 or Phe200 mutations. Position 194, showed the existence of 3 different variants carrying either Ile, Met or a polar Asn. Reference sequences of *T. trichiura*, and *T. suis* also show different variants at this position. *In silico*, a truncation was predicted in 3 of the 7 TBB genes as shown in SI Figure 5, which warrants further transcriptomic verification. To further investigate, if the variants at position 194 were reproducible in an independent Nanopore sequencing experiment of a single isolate, a region was amplified that was present in all genes and contained the predicted variants (primer sequences are provided in SI Figure 6). As shown in SI Figure 6, the different variants at position 194 were confirmed and the coverage ratio from the IGV software indicated that 8% of all reads carried an Asn, 37% carried an Ile, and 55% a Met, indicating that out of all orthologs of the TBB gene, one ortholog is present in the Asn form. Notably, these variants are very close to the predicted binding site of albendazole and are at variable sites compared to reference sequences.

**Figure 5:**
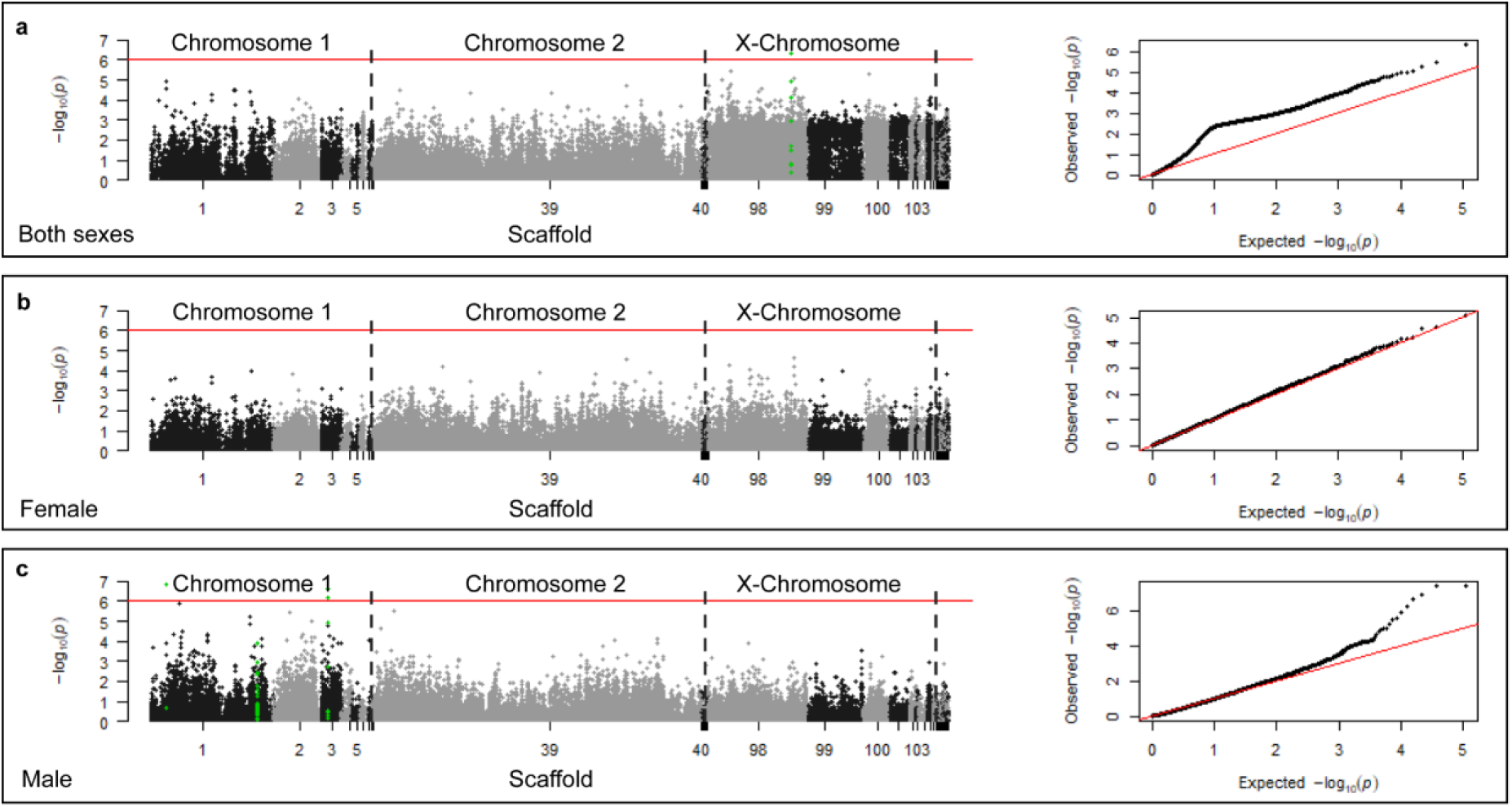
Genome wide association study of worms expelled after albendazole-ivermectin treatment (T1) versus worms expelled after oxantel-pamoate treatment (T2). GWAS displaying p-values of missense SNPs associated to treatment outcome (T1 vs T2) using a linear model without population stratification in PLINK. PCA is provided in the SI. A total of 24 GWAS’ were performed as described in the methodology section, the results of all of them are provided in the SI. The linear model without population stratification yielded the most hits while maintaining stable QQ plots for both sexes individually. Variants were filtered at a minor allele frequency of 0.05, a missing genotype frequency of 0.05, and a deviation from the Hardy-Weinberg Equilibrium at a significance level of p=0.001. The red line indicates the Bonferroni corrected significance of 8.89E-7 at α=0.05 and 56273 mutations. The scaffolds are separated by black/grey colors. The green dots indicate all SNPs found on the specific gene of interest carrying a SNP with genome wide significance. a Data of both sexes. 247 males, 471 females, and 3 ambiguous 542 cases as in drug non-sensitive and 179 controls. b Only female SNPs. 0 males, 471 females, 333 cases as in drug non-sensitive and 138 controls. C Only male SNPs. 247 males, 0 females, 208 cases as in drug non-sensitive and 39 controls. All GWAS hits, from each of the 24 analyses, the sequence, the best blast hit, the location and functional annotation, where possible, is provided in the SI Table 7 “hits_gwas_all.xlsx”

The gene tree of the orthologous group of TBB, OG0000671, shown in Figure 4c gives insight into the history of these duplication events. The gene tree shows a similar distance in number of substitutions per site from *T. suis* gene 4606 to *T. incognita n. sp.* gene 34, compared to the distance from *T. incognita n. sp.* gene 472 to *T. incognita n. sp.* gene 34. This indicates that interspecies genetic variation is similar to intraspecies genetic variation for at least 2 clades of TBB genes. Clade one, including gene 7506, 529, 472, 460, 463, and clade two, including gene 34. Genes 475, 476, and 5082 were not considered, as they were found to be fragmented.

We then sought to conduct an unbiased investigation of gene families which are as expanded or more expanded compared to the β-tubulin gene in *T. incognita n. sp.* and identified gene families associated with microtubule function and mitotic processes. Massively expanded gene families as in ≥7 copies per gene in *T. incognita n. sp.* and ≤4 in all other species were investigated and a list of gene families and their annotation is provided in the supporting information (SI Table 6). From 81 gene families, 68 could be associated to DNA transposable elements and retrotransposons, 6 remained unknown and without blast hits and 7 could be assigned a functionality. Apart from the TBB gene, other genes related to microtubule function and flexibility were found within these 7 assignable orthologous groups. Amongst these highly duplicated genes is the motor protein kinesin and MAP65/Ase1, which promotes microtubule flexibility and prevents microtubule severing by cross-linking.(50, 51) Finally, Condensin, Transcription factor (TF) IIIC, and a haspin like kinase, were identified and are involved in chromatid condensation, organization and cohesion respectively.

To extrapolate the TBB duplication events from the assembled genome of one single worm to the population, we leveraged the ratio of the average coverage of illumina reads of the amplified TBB region compared to the longest autosomal contig present in the genome of all 747 worms as shown in Figure 4b. Intriguingly, the gene ortholog count was reflected in the average overall coverage showing a significant difference between worm sex (p-value < 0.001) but not showing any significance with regard to treatment. On average 11 copies were present in female worms and 8 copies in male worms. To compare the coverage ratios against species, where only one ortholog was identified, the raw sequencing data of 26 *T. suis* worms isolated in Côte d’Ivoire was used as a comparison, where, intriguingly, only 1-2 copies of the TBB was present. Thus, one unique feature about *T. incognita n. sp.* is the high number of the TBB gene orthologs. The coverage ratios were further compared to coverage ratios from publicly available raw sequencing data that was downloaded for *T. suis* (PRJNA20815, PRJNA20816, PRJNA179528) and *T. trichiura* (PRJNA304165), the same region of TBB was used and the coverage ratio was calculated again from the amplified TBB region to the autosomal region indicating one ortholog of the TBB gene.

This work and the nomenclatural acts it contains for the new species have been registered in ZooBank: LSID: 34409100-BF94-4A0D-9088-D28215D231F6. Furthermore, a female holotype has been archived at the natural history museum in Basel under the number NMB- 79 a.

### Genome wide association study does not provide substantial evidence of adaptation to drug treatment

To investigate potential SNPs associated with resistance, a genome wide association study (GWAS) was conducted with sequencing data of 747 individual worms, 721 of which passed the filtering, comparing the populations isolated after T1 and T2 respectively comprising of 179 drug sensitive (T1) and 542 drug non-sensitive (T2) isolates. Investigation of the PCA presented in SI Figure 7, showed that worm-sex contributed the most to population stratification. As a result, the GWAS was conducted once for both sexes pooled, and for male and female worms separately. In the PCA of the subpopulation, significant population stratification was associated with the village of isolation. However, no association was found to T1 or T2 in the subpopulation, justifying the consideration of population stratification only on a worm sex level. Both a logistical and linear model was used to investigate 1254246 overall mutations and a subset of 56273 missense mutations on *in-silico* predicted genes and a genome wide significance level of 3.99E-8 and 8.89E-7 respectively using the Bonferroni correction at α=0.05 on the combined sample cohort of male and female worms and on each sex individually resulting in a total of 24 different runs presented in Figure 5 and SI Figures 8-15. QQ-plots showed consistent artificial inflation when using combined data of male and female worms, as seen in Figure 5a (right half of the panel). Furthermore, population stratification occasionally resulted in increased deviations from expected and observed values, for the probability of association to treatment outcome. When running only male or female isolates, the artificial inflation was mitigated (Figure 5b, 5c on the right). In total, 29 SNPs were identified with genome wide significance in all the 24 GWAS’ combined and are presented with their sequence in SI Table 7. Using a linear model without population stratification on missense mutations on coding genes (Figure 5), a total of 3 genes carried SNPs with genome wide significance in the male population: g207, a muscle M-line assembly protein unc-89 homolog; g1478, a FSA C domain containing protein; and g2562, an integrase catalytic domain-containing protein. The GWAS with a linear model without population stratification on all SNPs yielded 22 hits on 10 genes, including a voltage-gated chloride ion channel.. No further missense SNPs were identified with a genome wide significance. The QQ-plots were most consistent in the logistic model without population stratification as shown in SI Figure 10. Using this model, no SNPs with genome wide significance were identified.

SNPs on expanded gene families were also investigated. However, no genome wide significant SNPs were found in any of the 24 GWAS’. The strongest association of a missense SNP associated to drug treatment on the TBB gene was the Asn194Ile mutation with a p-value of p = 0.12 which is far below the genome wide significance level.

## Discussion

Our study presents an in-depth characterization of *T. incognita n. sp.* found to be less susceptible to albendazole-ivermectin treatment and morphologically indistinguishable to *T. trichiura*. A genome wide association study and the investigation of genes previously associated with resistance, did not conclusively show adaptation to drug pressure within the same species of *Trichuris*.

The existence of a species of *Trichuris*, phylogenetically distinct from the canonical human-infective species, has been proposed in recent studies involving amplicon sequencing by Rahman *et al.*,Venkatesan *et al*., and already in 2012 by Nissen *et al.*(11, 52, 53) Adopting the genetic species concept, our data confirm the existence of this species at the whole genome level, using the species tree constructed from 12434 orthologous groups, and 535 mitogenomes of 747 individually isolated and sequenced worms, all falling into the same clade. The whole-genome assembly metrics align with established *Trichuris* genomes, underscoring the robustness of our dataset.(15, 16, 32) However, structural variations might be missed, as the *T. muris* genome was used as a reference for scaffolding, similar to what was done in literature.(16) The prediction of genes in the publicly available *T. trichiura*, *T. suis*, *T. muris* and *Trichinella spiralis* genomes, and the newly assemble *T. incognita n. sp.* genome, using the same reference database, allowed the exploration of orthologous gene groups and duplication events across the species tree. The identification of a family of serpins and chemotryphsin-like serine proteases uniquely present in human infecting *Trichuris* may be of particular interest, as these proteins, which were identified to be upregulated in the stichosome of *T. suis*, have gained particular interest in the field of immunomodulatory diseases in general and vaccine development for *T. spiralis*.(15, 54, 55) Our investigation of duplication events across the species tree revealed a unique characteristic of *T. incognita n. sp.*: A high number of TBB gene orthologs. Duplication events are encountered across the tree of life and constitute an important mechanism to provide new genetic material. In eukaryotes, including *C. elegans*, gene duplication rates are estimated to be as frequent as single-nucleotide polymorphisms. However, they have never been investigated within the *Trichuris* genus.(56, 57) Drawing upon previous studies linking gene duplication to drug adaptation across diverse parasitic organisms, and deducing modes of action of drugs from observations of duplication events, we sought to investigate gene duplication events in connection to drug insensitivity.(58, 59) In total, 7 variants of the β-tubulin gene were identified in the whole genome, providing substantial redundancy that may allow for adaptation through sub-functionalization and increased gene expression. This contrasts with the low number of TBB orthologs in *T. trichiura*. It is of note that *T. trichiura* does not respond well to single benzimidazole treatment. Leveraging coverage ratios, duplication events could be confirmed in the whole population for each individual worm. Notably, while β- tubulin exhibited significant duplication, α-tubulin did not. Furthermore, massively expanded gene families are associated with microtubule dynamics and mitotic processes. Kinesin acts as intracellular shuttle and can de-polymerize microtubules. It has been shown to bind to negatively charged residues E410, D417 and E421 on the β-tubulin subunit.(60) MAP65/Ase1, a microtubule-associated protein is reported to promote microtubule flexibility and prevent microtubule severing by cross-linking.(50, 51) These findings suggest that the duplications observed are associated to repairing and protecting microtubule functionality which is intriguing considering the mode of action of albendazole, inhibiting microtubule formation.(61, 62) Further, highly duplicated genes including Condensin, Transcription factor (TF) IIIC and a haspin like kinase were associated to chromatid condensation, organization and cohesion respectively. While these genes have not been associated to benzimidazole resistance, they are linked to mitotic processes which in turn are connected to microtubule function with regards to the mitotic spindle.(63–65) *T. muris* is the only other organism with a high number of TBB orthologs and it has been shown that albendazole or mebendazole had an IC50 > 200 μg/mL in *T. muris*.(66) However, comparing the number of TBB orthologs of the population obtained after T1 to the population obtained after T2, showed no significant difference indicating that duplications are not the result of a recent adaptation nor an adaptation to drug pressure. Assuming an average mutation rate of 2 × 10^−8^ point mutations per gene per generation in *C. elegans* as an estimate, the duplication events would be dated back several centuries. These findings support the hypothesis that the observed lower cure rates in Côte d’Ivoire are not a result of resistance establishment, but potentially a result of different response rates of different *Trichuris* species. Finally, the functional significance of different TBB genes remains unknown and is subject for future studies.

Patterns of gene duplication, gain, and loss of function events in *T. incognita n. sp.* compared to within *T. suis* suggest a more ancient and divergent evolutionary lineage. Despite the close genetic relationship of *T. incognita n. sp.* to *T. suis*, our phylogenetic analysis demonstrates clear separation between *T. incognita n. sp.* and *T. suis* isolates from swine hosts in Côte d’Ivoire. While zoonosis events of trichuriasis from *T. suis* to *T. incognita n. sp.* have been reported,(67) these findings suggest that there may have been a zoonotic event sometime in the past between the human and swine host that was successful, but could not be observed currently. However, the occurrence of closely related *Trichuris* isolates from the colobus monkeys (*Colobus guereza kikuyensis*) raises the question, if these might belong to the same species of *Trichuris* and have zoonotic potential similar to *T. trichiura* species, which are found both within primates and humans.(32) Furthermore, J. Rivero *et al.* identified *Trichuris* species, which were closely related to the one found in the colobus monkey and may be the same species as *T. incognita n. sp.* based on ITS sequences or *cox*1 and *co*b genes.(43) If *T. incognita n. sp.* were able to infect non-human hosts, which is the case for *T. trichiura,* this would provide a refugia from drug pressure, which might prevent the establishment of anthelmintic resistance.(68)

With the unique experimental setup of the expulsion study, we were able to categorize helminths into drug sensitive and non-sensitive and thus enabling a genome wide association study (GWAS) to investigate the potential establishment of resistance to albendazole-ivermectin. MDA has been ongoing in Côte d’Ivoire for multiple decades,(4) which is well within the timeframe of a possible resistance establishment in STH through PC.(7) The most surprising finding of the GWAS may be, that there is no conclusive evidence to show a clear adaptation to drug treatment by comparing drug sensitive and non-sensitive *T. incognita* worms. Depending on the set of SNPs analyzed, regression model chosen for the GWAS, logistic vs linear, and the inclusion of principal components to account for population stratification or not, SNPs of genome wide significance with acceptable QQ-plots were only observed using a linear model in the male population without considering population stratification. Two hits are worth mentioning, as they can be found on genes that are to some extent in context of the mode of action of albendazole or ivermectin: a homolog of the muscle M-line assembly protein unc-89 found in *C. elegans* and a voltage gated chloride ion channel. The first one is a structural component of the muscle M line which regulates Ca^2+^ signaling and is involved in preventing the degradation of microtubule severing protein mei-1 by binding mel-26, which may be interesting regarding albendazoles mode of action.(69) The latter is interesting regarding the glutamate gated chloride ion channel binding activity of ivermectin, leading to increased hyperpolarization in the neuromuscular junction. (70) However, while these hits will be subject of future investigation, there is no clear indication of resistance establishment in the analysis conducted. This is in contrast to reported quantitative trait loci (QTL) associated with ivermectin resistance in *H. contortus* and *C. elegans* found on the cky- 1 gene,(14) and transmembrane proteins such as the chloride ion channel avr-15,(71) all of which did not show an association with genome wide significance in this study. Importantly, our experimental design also has limitations, such as the potential failure of capturing of multi-loci resistance mechanisms against a combination treatment, or gene expression related mechanisms, which might be visible in transcriptomic experiments.

The expulsion study also allowed us to gain insights into the fecundity and expulsion dynamics of *T. incognita*. Following the administration of albendazole and ivermectin during T1, only 271 out of 1098 worms were expelled, which is in-line with previously observed low cure rates in Côte d’Ivoire.(9, 72) The female to male ratio is comparable to recent reports for *T. trichiura*.(17) The fecundity of around 14,300 eggs per female per day of *T. incognita n. sp.* matches literature reported values of approximately 16,700 for *T. trichiura*.(17) Overall, these results suggest that the biology of *T. incognita n. sp.* in terms of fecundity is comparable to *T. trichiura*. Also morphologically, *T. incognita n. sp.* and *T. trichiura* were comparable. The presented morphological characteristics of *T. incognita n. sp.* do not enable a morphological distinction to *T. trichiura* indicating the limits of the morphological species concept for the *Trichuris* genus.(43) The morphological indistinguishability of *T. incognita n. sp.*, *T. trichiura* and *T. suis* worms may be the reason why this species has remained undescribed for many decades and emphasizes the need of diagnostic tools able to differentiate between these worms to gain further insight into the transmission dynamics and prevalence of this new species, as it responds differently to drug treatment. Intriguingly in a study from Rahman *et al*., analyzing a sample cohort from 2019 in similar regions, both *T. trichiura* and *T. incognita* were detected, where in our study exclusively *T. incognita* was observed in 2022.(53) Longitudinal studies investigating a potential connection between species distribution and drug efficacies will be of particular interest.

This study demonstrates that human trichuriasis can be caused by multiple species of whipworm, and that the differences in response rates may be a result of species responding differently to drug treatment, as opposed to an establishment of resistance. While the current WHO guideline for treatment of *T. trichiura* (combination therapy with (ALB/IVM) should still stand the current findings show the necessity to incorporate molecular diagnostics into clinical trials and to develop a new treatment against *T. incognita n. sp*.

## Acknowledgements

We would like to acknowledge Dr. John Gilleard and Dr. Abhinaya Venkatesan from the Faculty of Veterinary Medicine, Host-Parasite Interactions Program, University of Calgary, AB T2N1N4, Canada for the fruitful discussions on *Trichuris* phylogeny. We are grateful to the European Research Council for financial support (Nr. 101019223).

## Supporting Information Figures

**SI Figure 1:**
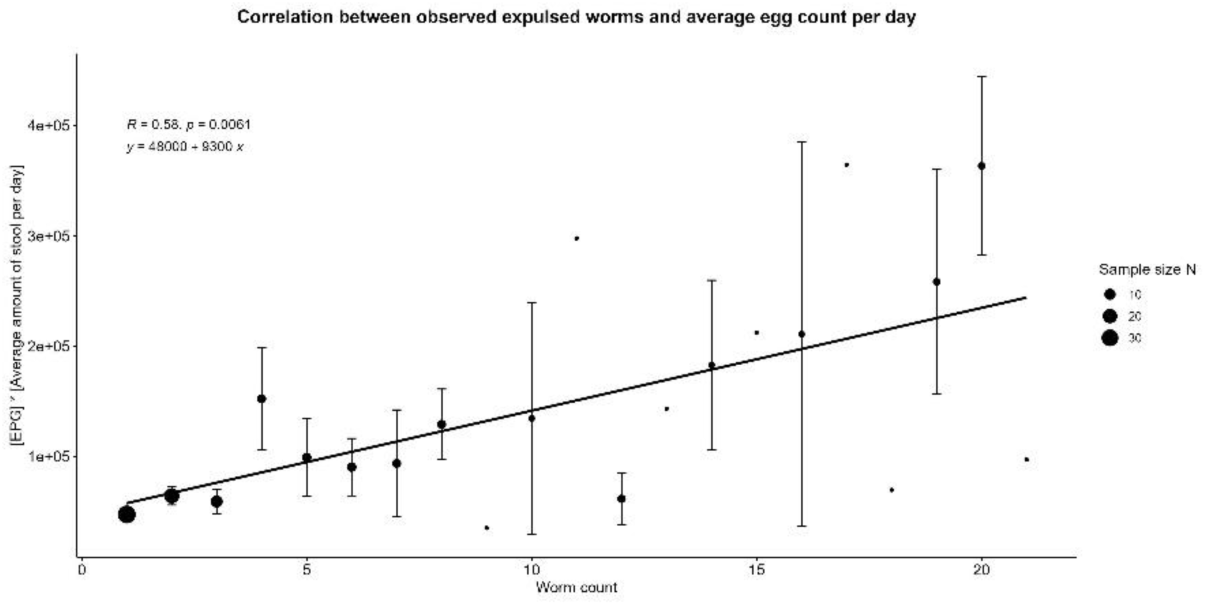
Correlation between observed expelled worms and average egg count per day. Error bars indicate standard error with no error bar present if there was only one value.

**SI Figure 2:**
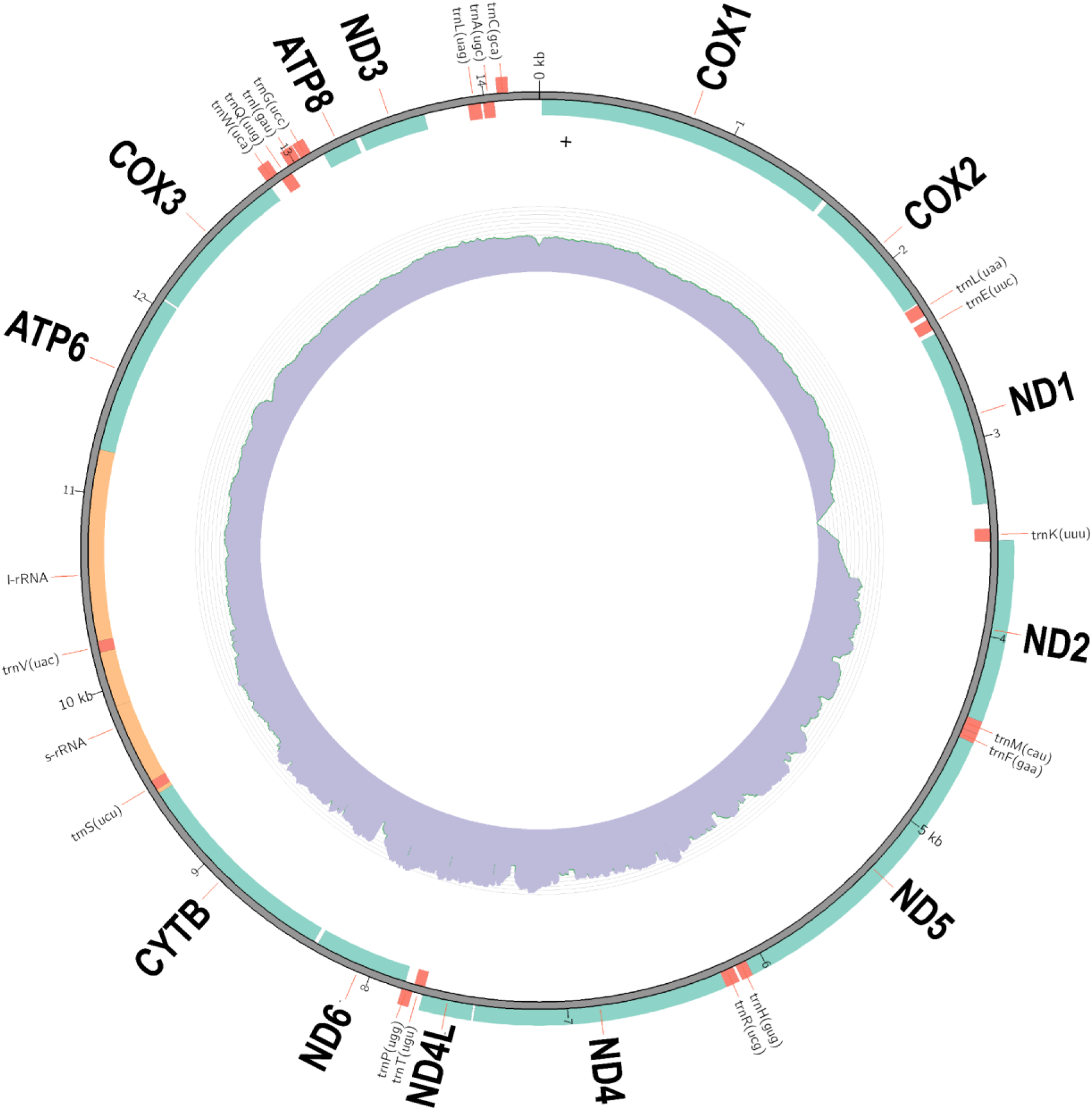
Representative image of a reconstructed mitochondrial genome containing 13 protein coding genes: *cox*1-3, *nad*1-6, *nad*4L, *atp*6, *atp*8 and *co*B. Turquois represents the coding sequence of genes, the inner ring represents the normalized coverage, light orang represents s- and l-rRNA

**SI Figure 3:**
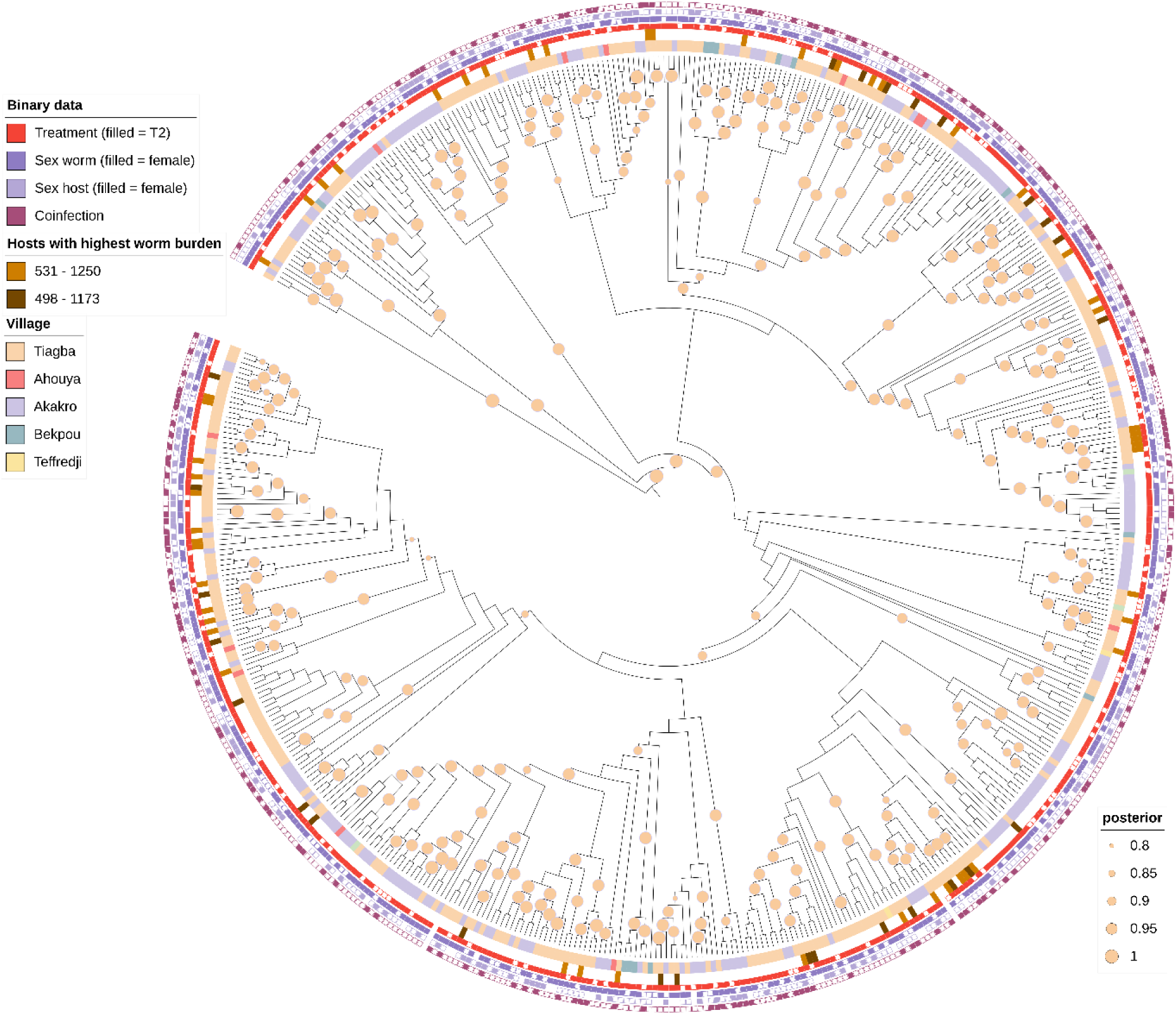
Phylogenetic tree based on mitochondrial sequences from 535 isolated *T. incognita n. sp.* worms from human hosts. The circle on branches indicates the posterior distribution. The outmost 4 rings represent binary metadata. A filled red tile indicates that the worm was isolated after T2 where the empty tile indicates isolation after T1. The dark violet tile indicates the worm sex and the light violet tile the host sex while purple indicates if a coinfection with other worms was observed. The middle ring indicates two hosts: A host with the ID number “531 – 1250” in light brown and a host with the ID number “498 – 1173” in dark brown, showing that the worms isolated from the two different hosts are scattered throughout the phylogenetic tree. The inner circle indicates the village from which the worm was isolated.

**SI Figure 4:**
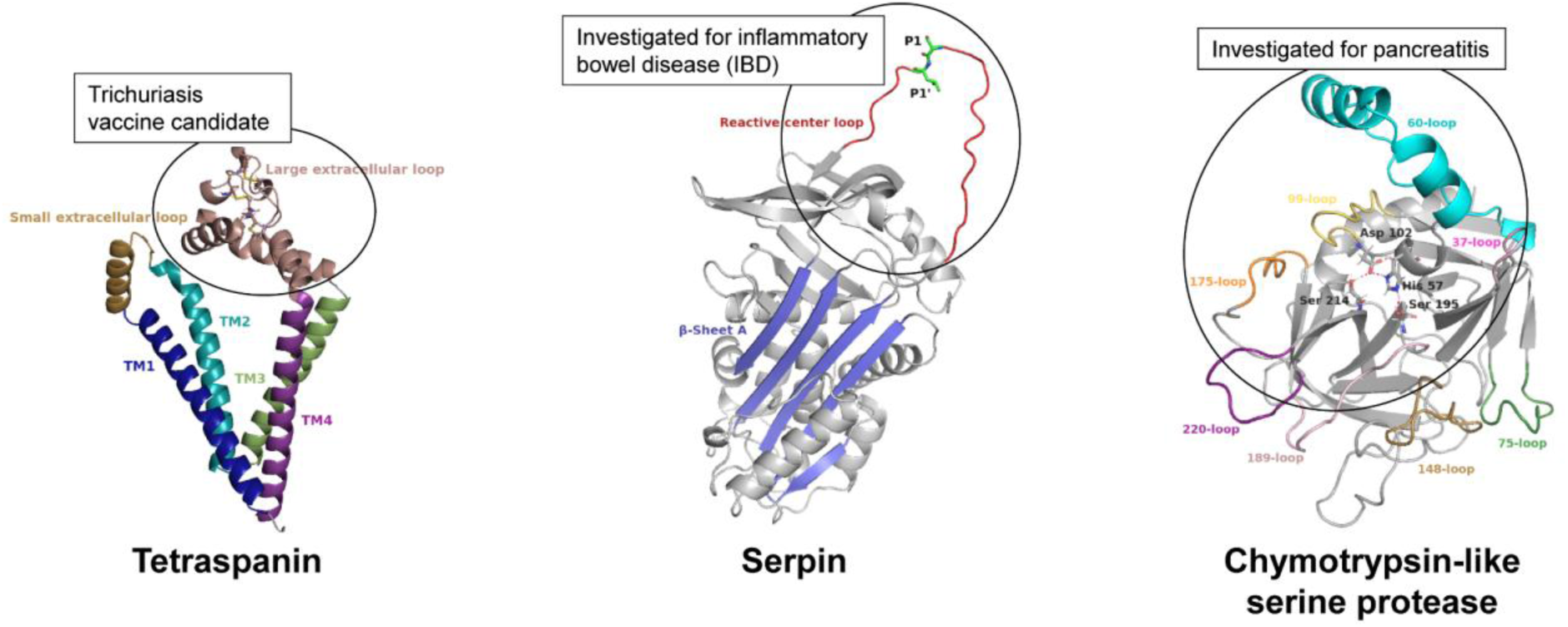
Tetraspanin, Serpin and Chymotrypsin-like serine protease from analyzing orthologous groups in Venn diagram sections shared by *T. incognita n. sp.*, *T. trichiura* (Serpin and Chymotrypsin-like serine protease) and *T. muris* (Tetraspanin).

**SI Figure 5:**
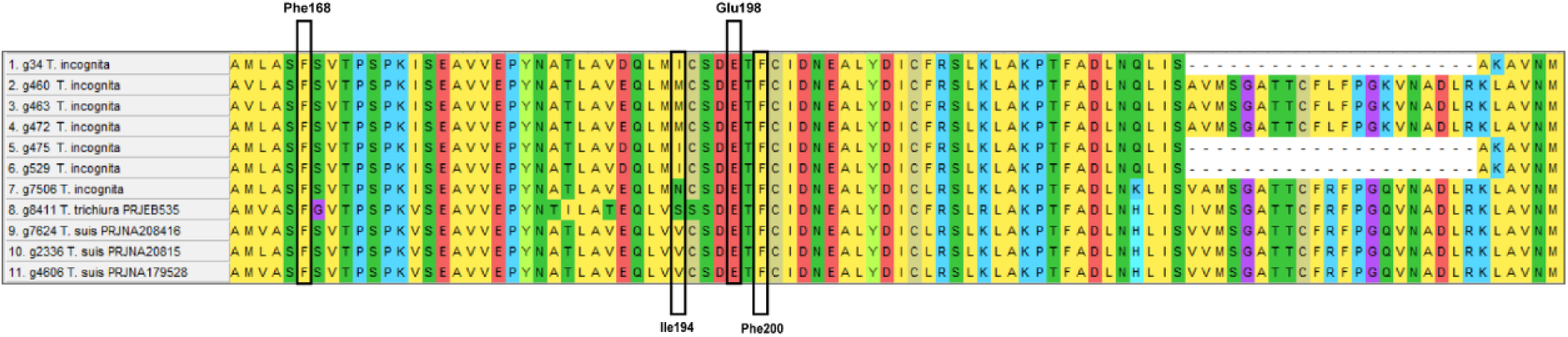
Beta tubulin genes found in *T. incognita n. sp.* compared to *T. suis* and *T. trichiura*, predicted from the braker2 pipeline. Black indicates positions carrying known mutations (Phe168, Glu198, Phe200), observed variable position (Ile194). A truncation was predicted *in silico* on 3 of the 7 genes. Gene IDs from top to bottom: g34, g460, g463, g472, g475, g529 and g7506.

**SI Figure 6:**
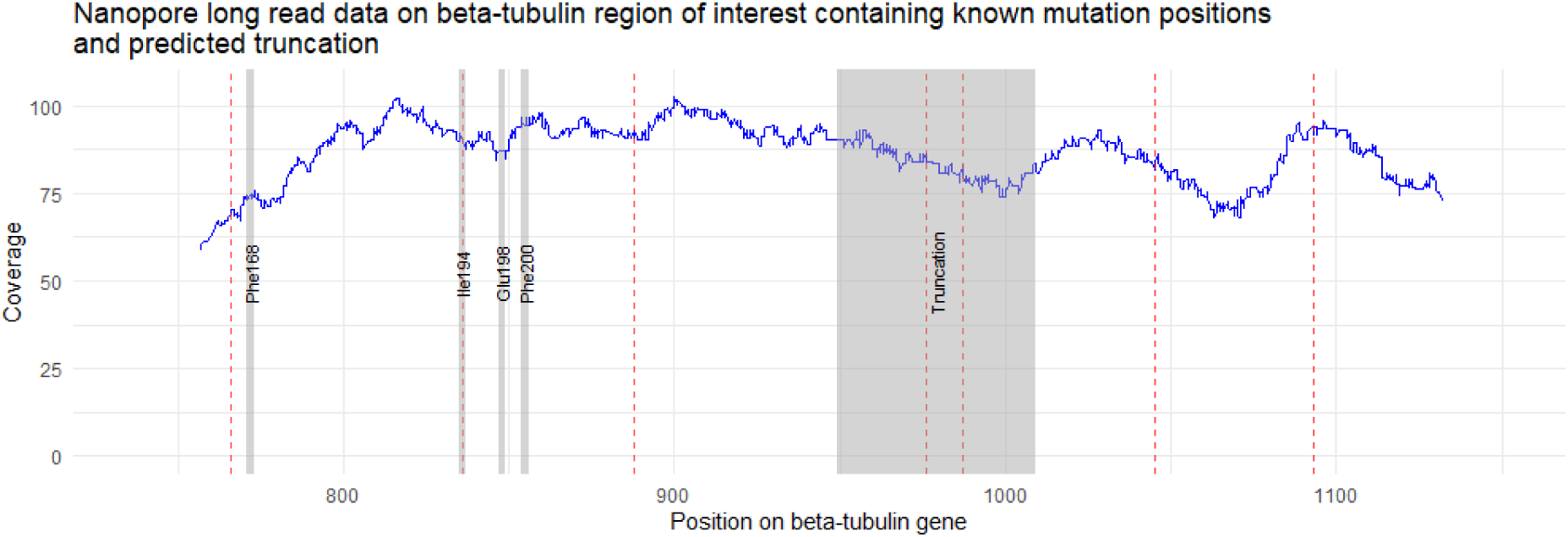
Coverage and variants from an independent PCR and Nanopore genome sequencing experiment of the regions containing known mutations of TBB and the *in silico* predicted truncation. Blue line indicates coverage, red lines the called variants using bcftools. Gray indicating positions carrying known mutations (Phe168, Glu198, Phe200), observed variable position (Ile194) and predicted truncation indicating that the truncation was not observed from the amplified genomic sequence. Forward primer sequence: AAAGAGACCGGACATTTCGC. Reversed primer sequence: TGAATTGCCTGGTTTCTAGAATG.

**SI Figure 7:**
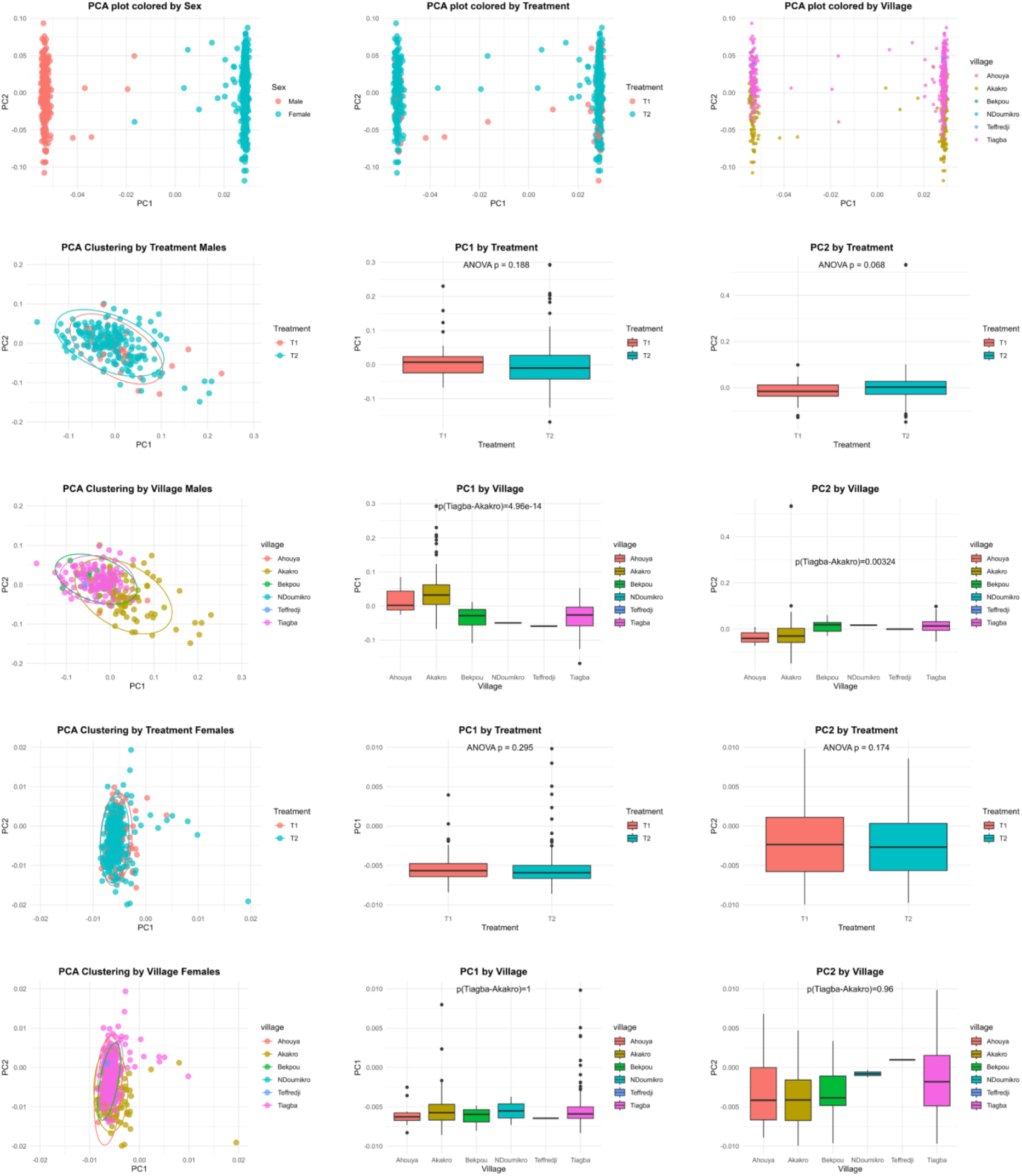
Summary of the PCA of all missense mutations on coding genes for the GWAS showing no population structure associated to treatment. a-c PCA plots of all missense mutations showing that worm sex has the biggest impact on population stratification. d-i PCA plots of only male worms showed significant population stratification by Tukey multiple comparisons of means in PC1 between the villages Tiagba-Akakro (p=4.96 E-14) and Bekpou-Akakro (p=2.79 E-05) and in PC2 between the villages Tiagba-Akakro (p=0.0032). No association to the treatment (ANOVA p > 0.05) was found in PC1 or PC2. j-o PCA plots of only female worms showed significant population stratification by Tukey multiple comparisons of means in PC1 between the villages Tiagba-Ahouya (p=0.0009) and Ahouya-Akakro (p=0.0006) and in PC2 between the villages Ahouya-Akakro (p=0.027). No association to the treatment (ANOVA p > 0.05) was found in PC1 or PC2.

**SI Figure 8:**
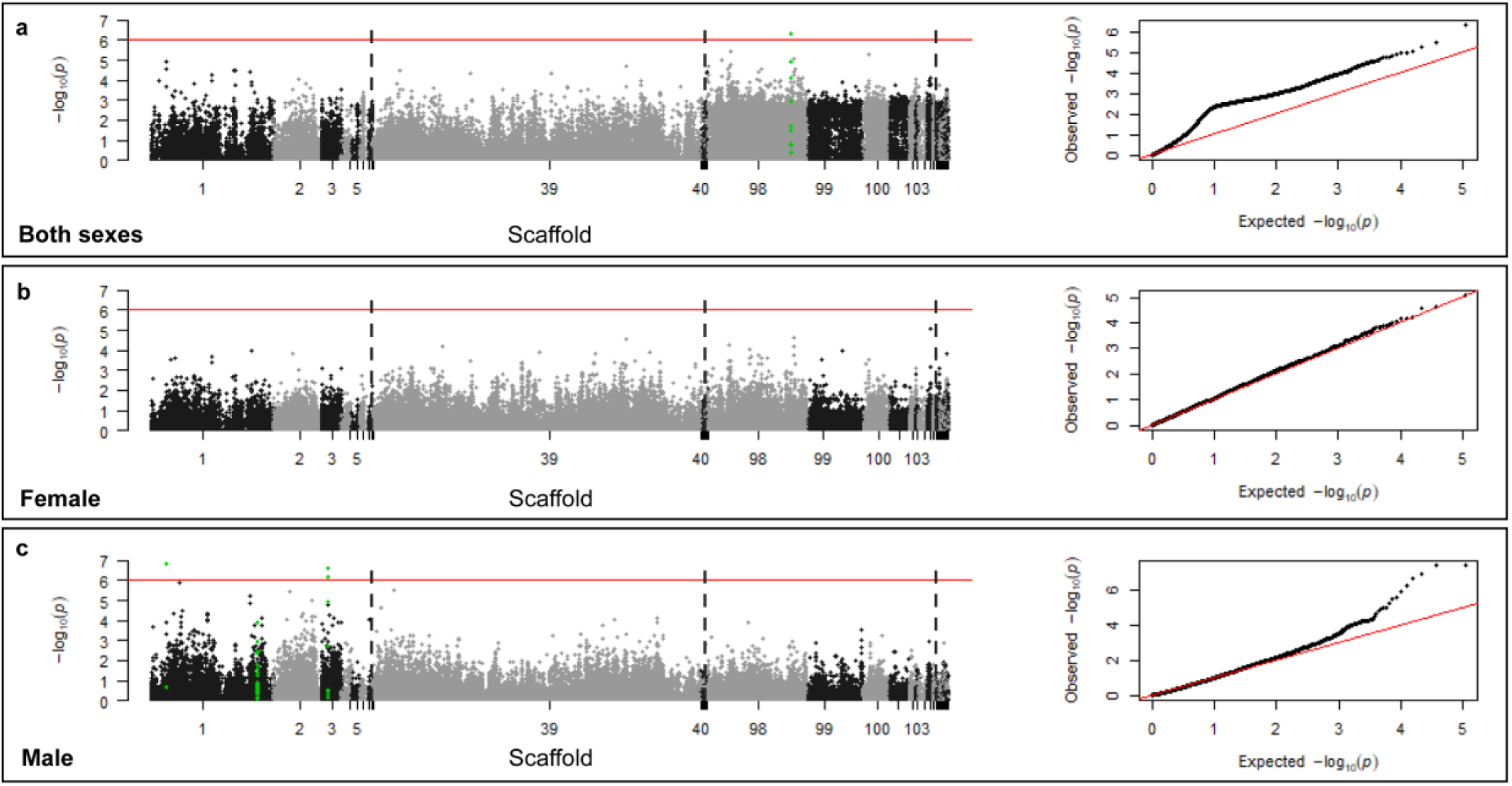
GWAS displaying p-values of missense SNPs on predicted genes associated to treatment outcome (T1 vs T1) using a linear model without population stratification in PLINK. Variants were filtered at a minor allele frequency of 0.05, a missing genotype frequency of 0.05, and a deviation from the Hardy-Weinberg Equilibrium at a significance level of p=0.001. The red line indicates the Bonferroni corrected significance of 8.89E-7 at α=0.05 and 56273 mutations. The scaffolds are separated by black/grey colors. The green dots indicate all SNPs found on the specific gene of interest carrying a SNP with genome wide significance. **a** Data of both sexes. 247 males, 471 females, and 3 ambiguous 542 cases as in drug non-sensitive and 179 controls. **b** Only female SNPs. 0 males, 471 females, 333 cases as in drug non-sensitive and 138 controls. **c** Only male SNPs. 247 males, 0 females, 208 cases as in drug non-sensitive and 39 controls. All GWAS hits, from each analysis, the sequence, the best blast hit, the location and functional annotation, where possible, is provided in the SI Table 7 “hits_gwas_all.xlsx”

**SI Figure 9:**
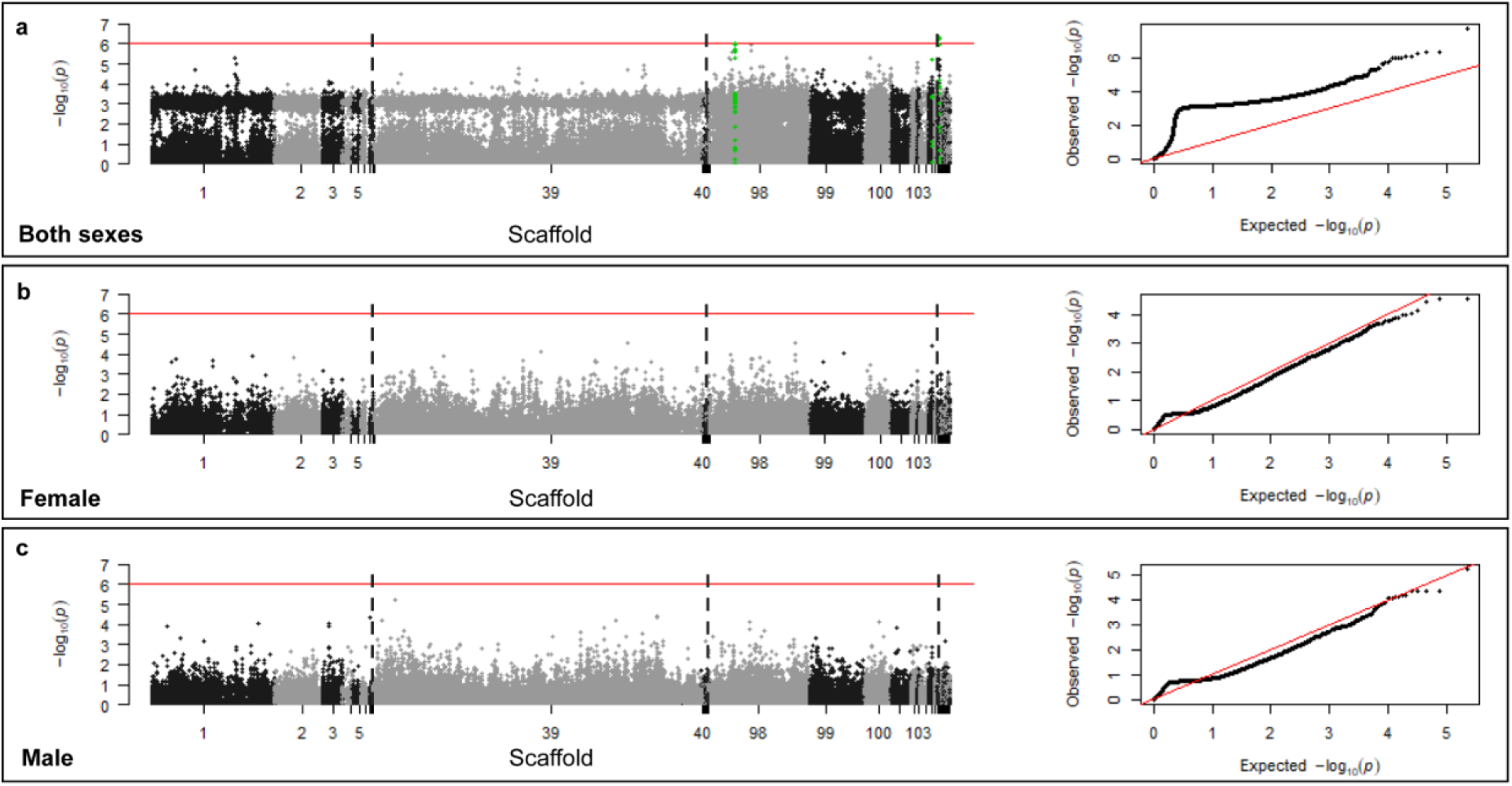
GWAS displaying p-values of missense SNPs on predicted genes associated to treatment outcome (T1 vs T1) using a linear model with population stratification in PLINK. Variants were filtered at a minor allele frequency of 0.05, a missing genotype frequency of 0.05, and a deviation from the Hardy-Weinberg Equilibrium at a significance level of p=0.001. The red line indicates the Bonferroni corrected significance of 8.89E-7 at α=0.05 and 56273 mutations. The scaffolds are separated by black/grey colors. The green dots indicate all SNPs found on the specific gene of interest carrying a SNP with genome wide significance. **a** Data of both sexes. 247 males, 471 females, and 3 ambiguous 542 cases as in drug non-sensitive and 179 controls. **b** Only female SNPs. 0 males, 471 females, 333 cases as in drug non-sensitive and 138 controls. **c** Only male SNPs. 247 males, 0 females, 208 cases as in drug non-sensitive and 39 controls. All GWAS hits, from each analysis, the sequence, the best blast hit, the location and functional annotation, where possible, is provided in the SI Table 7 “hits_gwas_all.xlsx”

**SI Figure 10:**
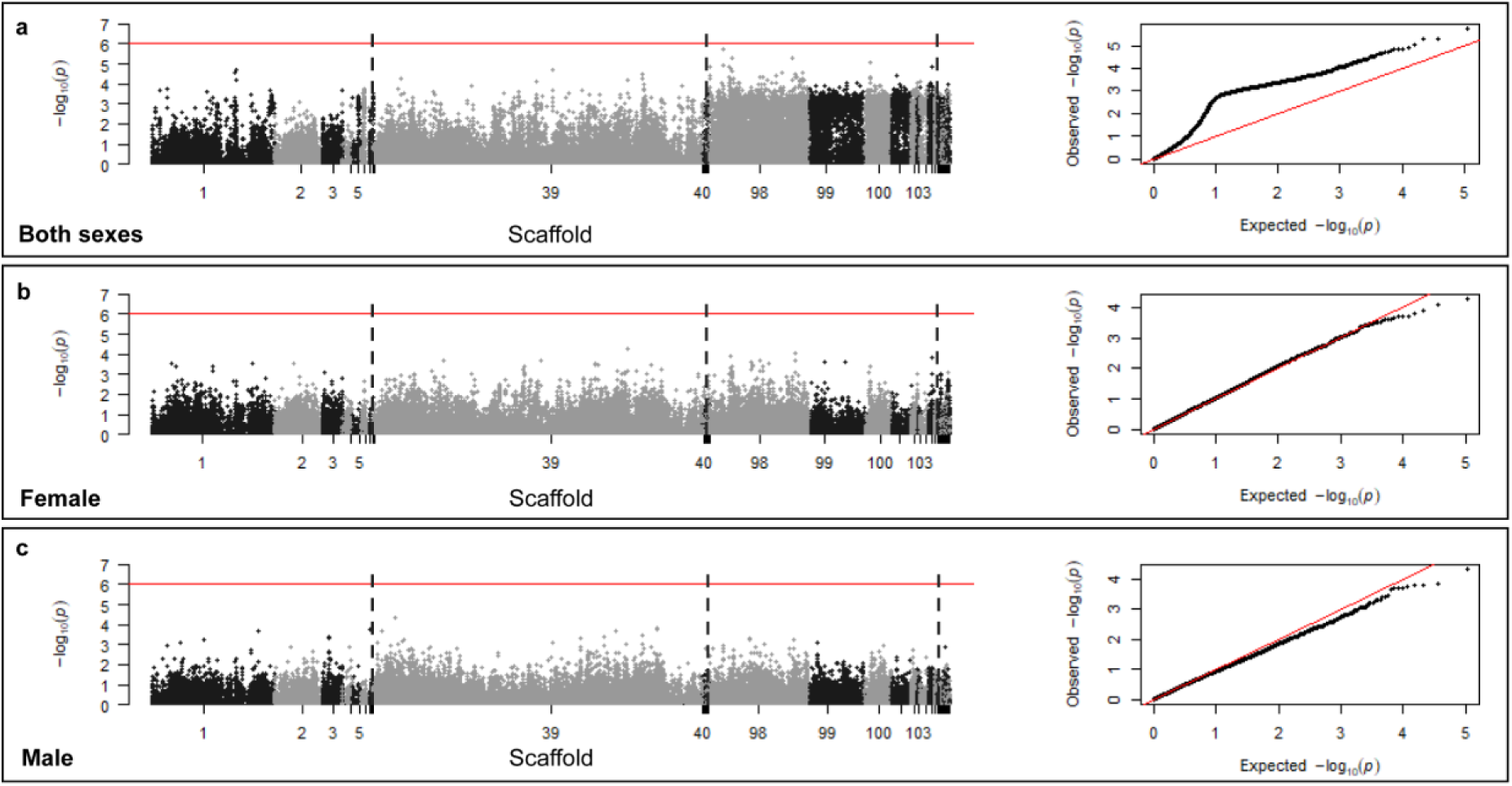
GWAS displaying p-values of missense SNPs on predicted genes associated to treatment outcome (T1 vs T1) using a logistic model without population stratification in PLINK. Variants were filtered at a minor allele frequency of 0.05, a missing genotype frequency of 0.05, and a deviation from the Hardy-Weinberg Equilibrium at a significance level of p=0.001. The red line indicates the Bonferroni corrected significance of 8.89E-7 at α=0.05 and 56273 mutations. The scaffolds are separated by black/grey colors. The green dots indicate all SNPs found on the specific gene of interest carrying a SNP with genome wide significance. **a** Data of both sexes. 247 males, 471 females, and 3 ambiguous 542 cases as in drug non-sensitive and 179 controls. **b** Only female SNPs. 0 males, 471 females, 333 cases as in drug non-sensitive and 138 controls. **c** Only male SNPs. 247 males, 0 females, 208 cases as in drug non-sensitive and 39 controls. All GWAS hits, from each analysis, the sequence, the best blast hit, the location and functional annotation, where possible, is provided in the SI Table 7 “hits_gwas_all.xlsx”

**SI Figure 11:**
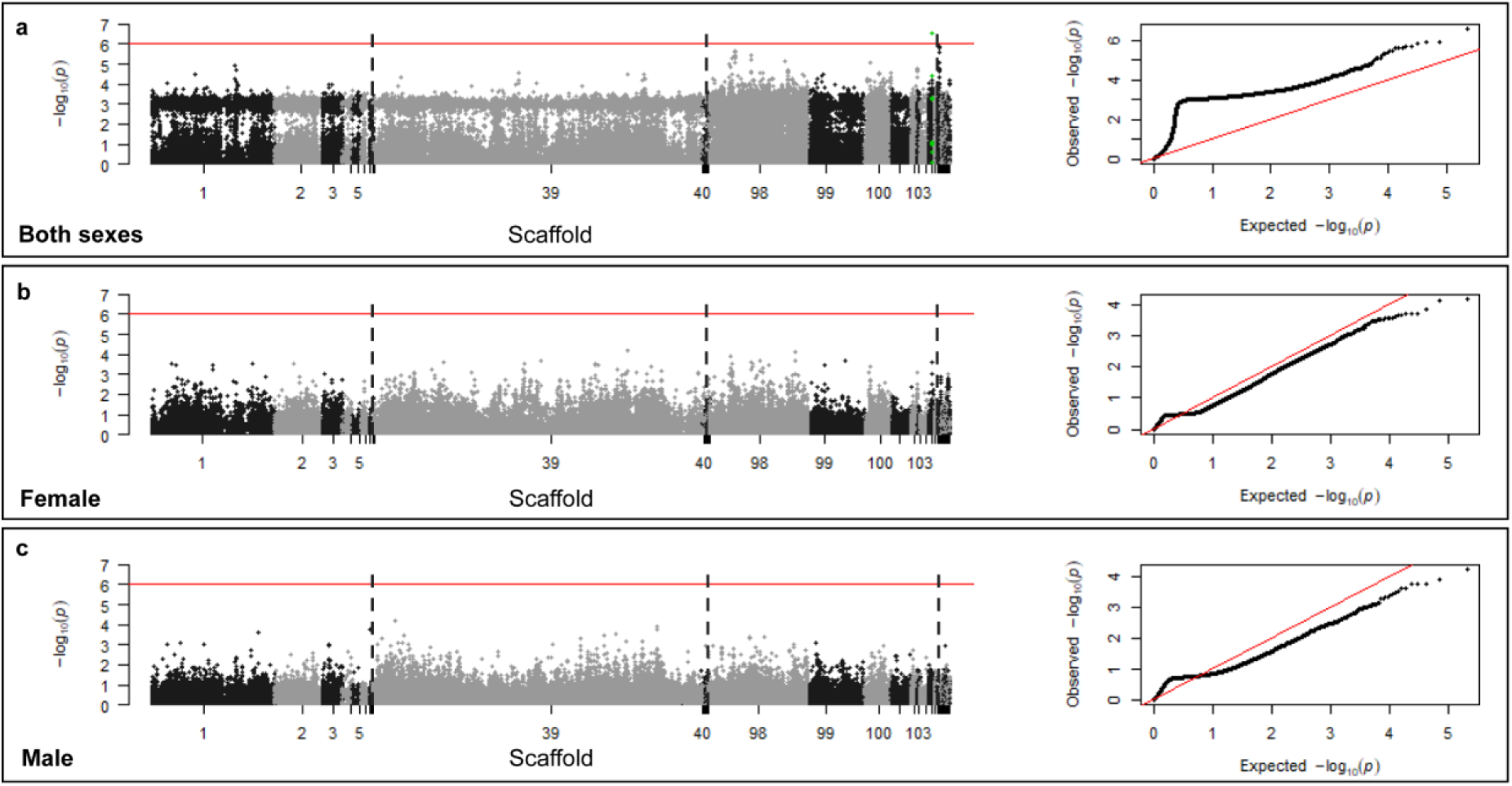
GWAS displaying p-values of missense SNPs on predicted genes associated to treatment outcome (T1 vs T1) using a logistic model with population stratification in PLINK. Variants were filtered at a minor allele frequency of 0.05, a missing genotype frequency of 0.05, and a deviation from the Hardy-Weinberg Equilibrium at a significance level of p=0.001. The red line indicates the Bonferroni corrected significance of 8.89E-7 at α=0.05 and 56273 mutations. The scaffolds are separated by black/grey colors. The green dots indicate all SNPs found on the specific gene of interest carrying a SNP with genome wide significance. **a** Data of both sexes. 247 males, 471 females, and 3 ambiguous 542 cases as in drug non-sensitive and 179 controls. **b** Only female SNPs. 0 males, 471 females, 333 cases as in drug non-sensitive and 138 controls. **c** Only male SNPs. 247 males, 0 females, 208 cases as in drug non-sensitive and 39 controls. All GWAS hits, from each analysis, the sequence, the best blast hit, the location and functional annotation, where possible, is provided in the SI Table 7 “hits_gwas_all.xlsx”

**SI Figure 12:**
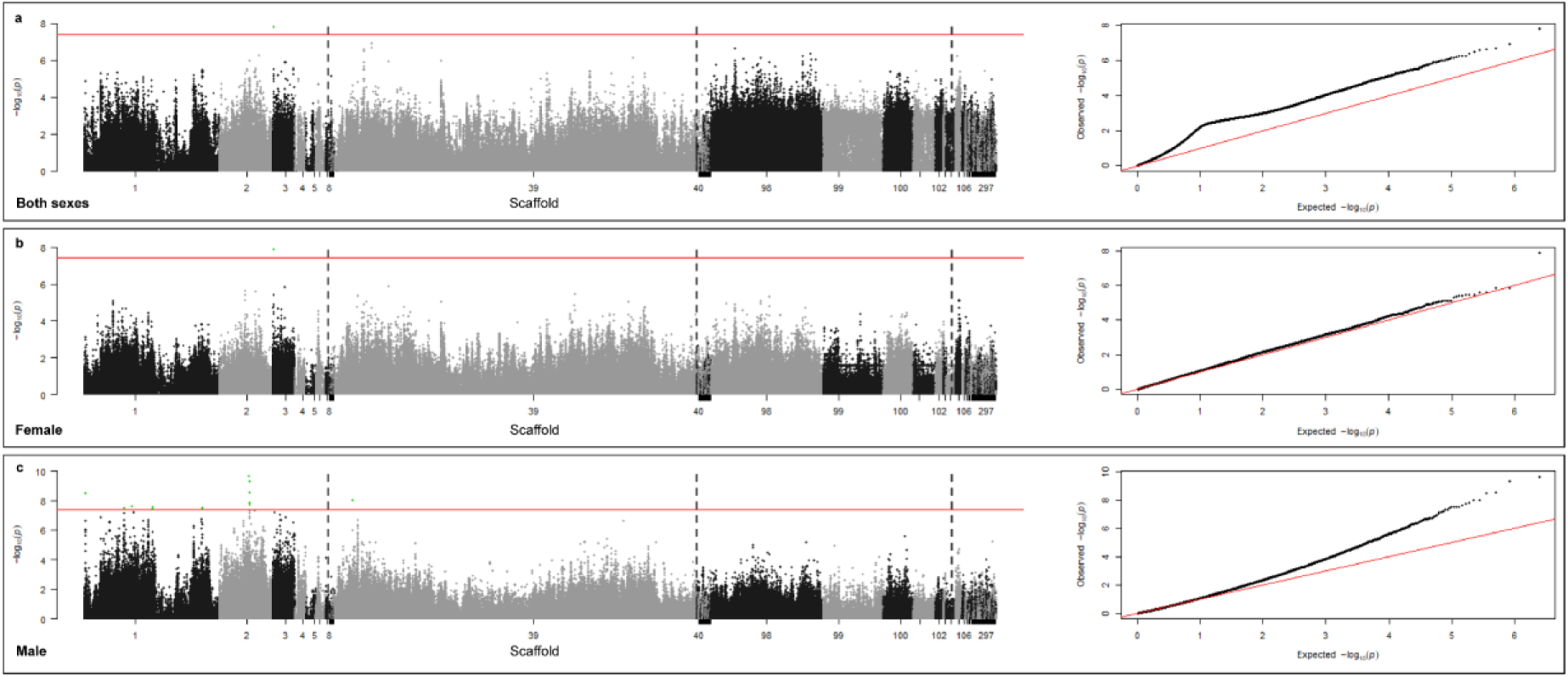
GWAS displaying p-values of all SNPs associated to treatment outcome (T1 vs T1) using a linear model without population stratification in PLINK. Variants were filtered at a minor allele frequency of 0.05, a missing genotype frequency of 0.05, and a deviation from the Hardy-Weinberg Equilibrium at a significance level of p=0.001. The red line indicates the Bonferroni corrected significance of 3.99 E-8 at α=0.05 and 1254247 mutations. The scaffolds are separated by black/grey colors. The green dots indicate all SNPs found on the specific gene of interest carrying a SNP with genome wide significance. **a** Data of both sexes. 247 males, 471 females, and 3 ambiguous 542 cases as in drug non-sensitive and 179 controls. **b** Only female SNPs. 0 males, 471 females, 333 cases as in drug non-sensitive and 138 controls. **c** Only male SNPs. 247 males, 0 females, 208 cases as in drug non-sensitive and 39 controls. All GWAS hits, from each analysis, the sequence, the best blast hit, the location and functional annotation, where possible, is provided in the SI Table 7 “hits_gwas_all.xlsx”

**SI Figure 13:**
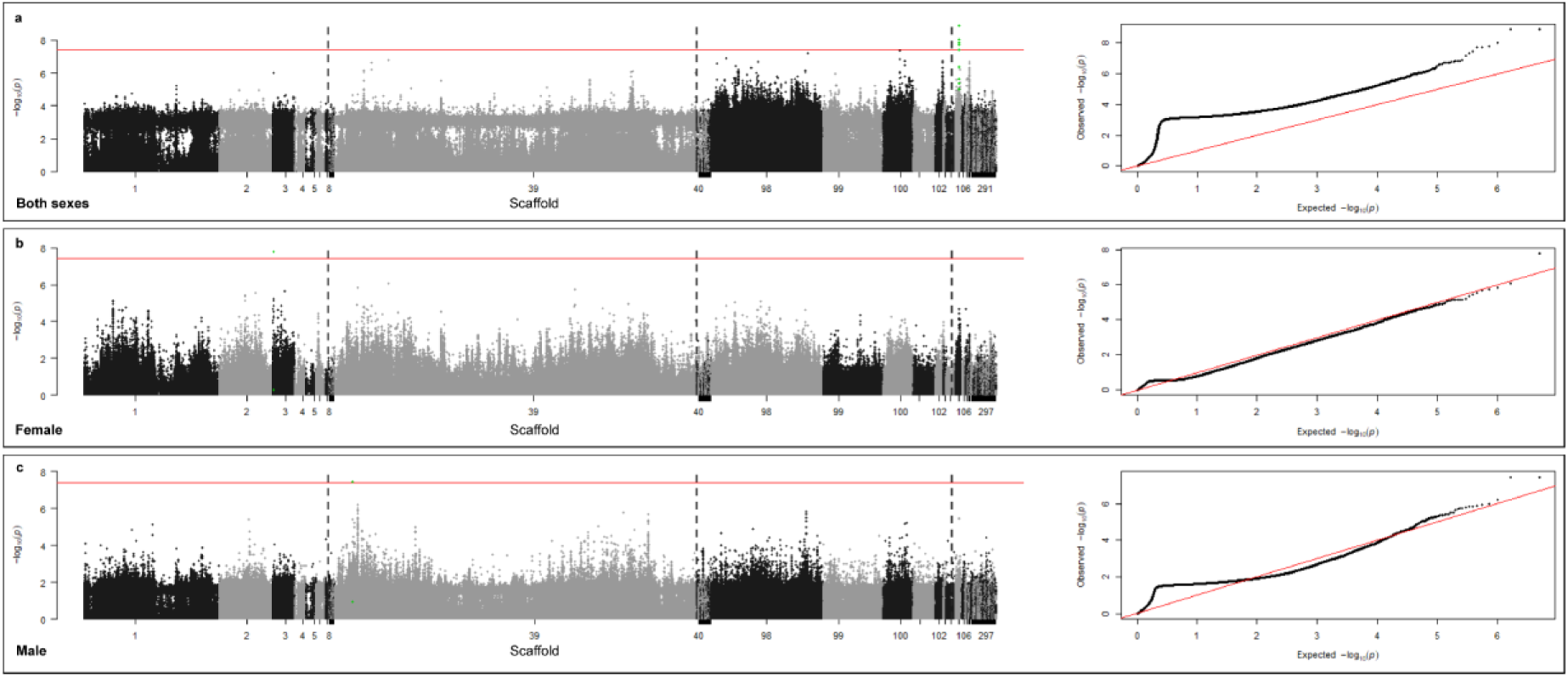
GWAS displaying p-values of all SNPs associated to treatment outcome (T1 vs T1) using a linear model with population stratification in PLINK. Variants were filtered at a minor allele frequency of 0.05, a missing genotype frequency of 0.05, and a deviation from the Hardy-Weinberg Equilibrium at a significance level of p=0.001. The red line indicates the Bonferroni corrected significance of 3.99 E-8 at α=0.05 and 1254247 mutations. The scaffolds are separated by black/grey colors. The green dots indicate all SNPs found on the specific gene of interest carrying a SNP with genome wide significance. **a** Data of both sexes. 247 males, 471 females, and 3 ambiguous 542 cases as in drug non-sensitive and 179 controls. **b** Only female SNPs. 0 males, 471 females, 333 cases as in drug non-sensitive and 138 controls. **c** Only male SNPs. 247 males, 0 females, 208 cases as in drug non-sensitive and 39 controls. All GWAS hits, from each analysis, the sequence, the best blast hit, the location and functional annotation, where possible, is provided in the SI Table 7 “hits_gwas_all.xlsx”

**SI Figure 14:**
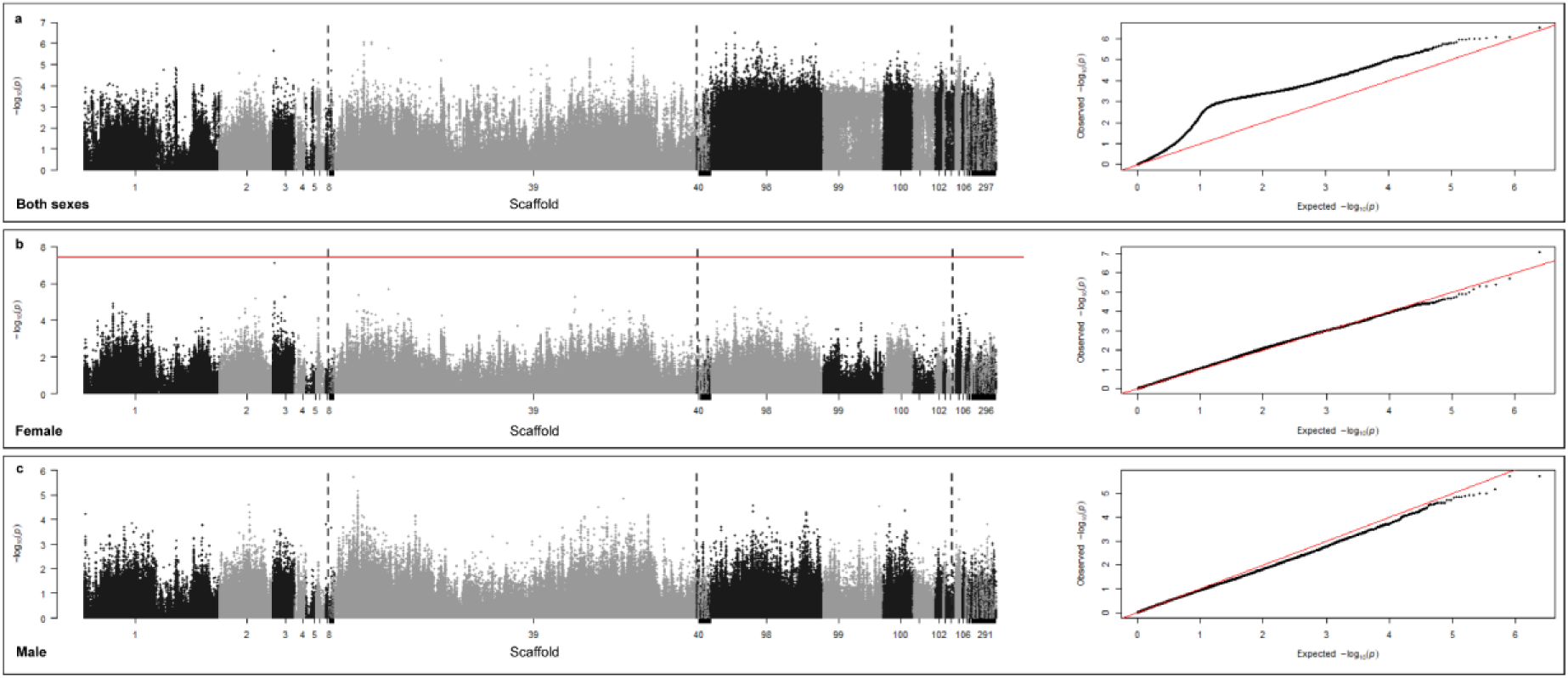
GWAS displaying p-values of all SNPs associated to treatment outcome (T1 vs T1) using a logistic model without population stratification in PLINK. Variants were filtered at a minor allele frequency of 0.05, a missing genotype frequency of 0.05, and a deviation from the Hardy-Weinberg Equilibrium at a significance level of p=0.001. The red line indicates the Bonferroni corrected significance of 3.99 E-8 at α=0.05 and 1254247 mutations. The scaffolds are separated by black/grey colors. The green dots indicate all SNPs found on the specific gene of interest carrying a SNP with genome wide significance. **a** Data of both sexes. 247 males, 471 females, and 3 ambiguous 542 cases as in drug non-sensitive and 179 controls. **b** Only female SNPs. 0 males, 471 females, 333 cases as in drug non-sensitive and 138 controls. **c** Only male SNPs. 247 males, 0 females, 208 cases as in drug non-sensitive and 39 controls. All GWAS hits, from each analysis, the sequence, the best blast hit, the location and functional annotation, where possible, is provided in the SI Table 7 “hits_gwas_all.xlsx”

**SI Figure 15:**
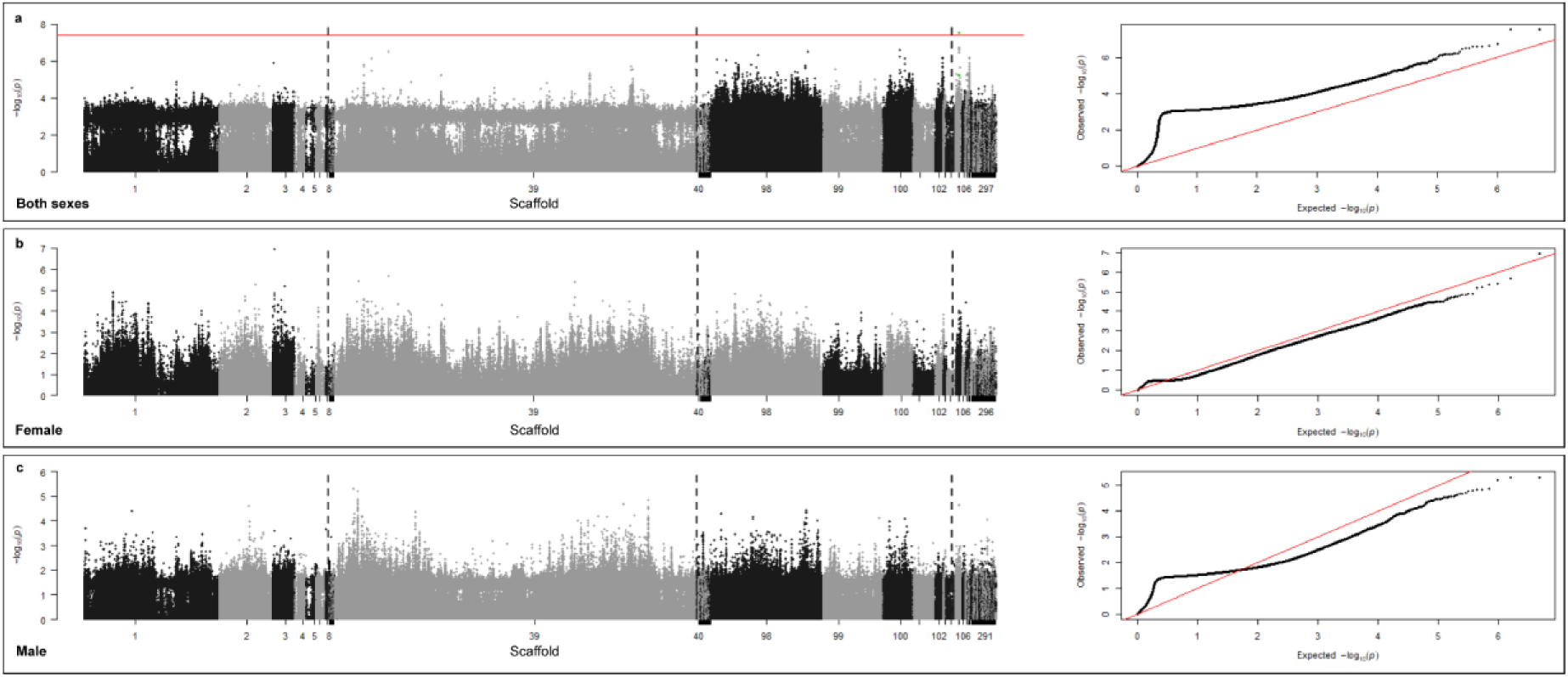
GWAS displaying p-values of all SNPs associated to treatment outcome (T1 vs T1) using a logistic model with population stratification in PLINK. Variants were filtered at a minor allele frequency of 0.05, a missing genotype frequency of 0.05, and a deviation from the Hardy-Weinberg Equilibrium at a significance level of p=0.001. The red line indicates the Bonferroni corrected significance of 3.99 E-8 at α=0.05 and 1254247 mutations. The scaffolds are separated by black/grey colors. The green dots indicate all SNPs found on the specific gene of interest carrying a SNP with genome wide significance. **a** and **b** Data of both sexes. 247 males, 471 females, and 3 ambiguous 542 cases as in drug non-sensitive and 179 controls. **c** and **d** Only female SNPs. 0 males, 471 females, 333 cases as in drug non-sensitive and 138 controls. **e** and **f** Only male SNPs. 247 males, 0 females, 208 cases as in drug non-sensitive and 39 controls. All GWAS hits, from each analysis, the sequence, the best blast hit, the location and functional annotation, where possible, is provided in the SI Table 7 “hits_gwas_all.xlsx”

**SI Table 1:** Five randomly sampled male and female worms (top) and literature reported values (bottom). Total length, posterior length, posterior width, anterior length and anterior width are provided, including the geometrical mean and standard deviation (top) or Min-Max values (bottom).

| Sample ID | Sex | Total length | Posterior length | Posterior width | Anterior length | Anterior width |
| --- | --- | --- | --- | --- | --- | --- |
| 995 | F | 36 | 12 | 1.2 | 24 | 0.09 |
| 787 | F | 48 | 15 | 0.8 | 33 | 0.15 |
| 66 | F | 44 | 13 | 0.6 | 31 | 0.12 |
| 618 | F | 45 | 11 | 0.7 | 34 | 0.15 |
| 368 | F | 40 | 12 | 0.6 | 28 | 0.11 |
| **Mean** |  | **42.39** | **12.53** | **0.75** | **29.77** | **0.12** |
| **SD** |  | **4.67** | **1.52** | **0.25** | **4.06** | **0.03** |
| Sample ID | Sex | Total length | Posterior length | Posterior width | Anterior length | Anterior width |
| 9 | M | 36 | 11 | 1 | 25 | 0.2 |
| 965 | M | 40 | 17 | 1.1 | 23 | 0.25 |
| 960 | M | 41 | 14 | 0.5 | 27 | 0.15 |
| 271 | M | 38 | 13 | 0.6 | 25 | 0.07 |
| 17 | M | 41 | 18 | 0.5 | 23 | 0.1 |
| **Mean** |  | **39.15** | **14.37** | **0.70** | **24.55** | **0.14** |
| **SD** |  | **2.17** | **2.88** | **0.29** | **1.67** | **0.07** |

|  | Total length (Min-Max) | Posterior length (Min-Max) | Posterior width (Min-Max) | Anterior length (Min-Max) | Anterior width (Min-Max) |
| --- | --- | --- | --- | --- | --- |
| **Females** |  |  |  |  |  |
| *T. incognita* n. sp. (Present study) | 36-45 | 11-15 | 0.6-1.2 | 24-34 | 0.09-0.15 |
| *T. trichiura* from J. Rivero *et al.* | 20-49 | 6-16 | 0.38-0.9 | 13-33 | 0.09-0.2 |
| **Males** |  |  |  |  |  |
| *T. incognita* n. sp. (Present study) | 36-41 | 11-15 | 0.5-1 | 23-27 | 0.07-0.25 |
| *T. trichiura* from J. Rivero *et al.* | 25-49 | 7-20 | 0.4-0.7 | 18-29 | 0.09-0.31 |

**SI Table 2.** Venn diagram of orthologous groups from predicted transcripts, where 175 were exclusively shared between T. incognita n. sp. and T. trichiura and their GO enrichment analysis. Availability of a *Trichuris* genome, closely related to *T. suis* but from a species naturally infecting humans, poses a unique opportunity to conduct an exploratory analysis of genes that might be crucial for the adaptation to a human host. We looked into the occurrence of orthologous groups throughout the genomes of *T. incognita n. sp.*, *T. suis*, *T. trichiura* and *T. muris* as shown in Table 2. Leveraging the close genetic relationship with *T. suis*, predicted transcripts were investigated that are shared with *T. trichiura* but do not occur in *T. suis* or *T. muris*. A GO term enrichment analysis was performed on this subset of genes using the predicted genes in *T. incognita n. sp.* as a background gene set and the full list of significant terms is provided in SI Table 3. A list of GO enriched terms provided in Table 2 indicate extracellular space to be the most significantly enriched term in the category of cellular components along with the nucleosom. Nucleic acid binding, tryptophan-tRNA ligase activity, structural constituent of chromatin and RNA binding were the most significant in molecular functions and finally tryptophanyl-tRNA aminoacylation, [2Fe-2S] cluster assembly and protein complex oligomerization in biological process. Finally, considering that the anterior section is in a most intimate relationship with the host, we searched for genes within this gene-set which Jex *et. al* identified to be upregulated in the stichosome and potentially have immunomodulatory effects like galactins, serpins, venom allergen–like proteins, apyrase or calreticulin and chymotrypsin-like serine proteases.(15) We identified one family of each serpins (OG0000305) and chymotrypsin-like serine proteases (OG0010319) that are shared only by *T. trichiura* and *T. incognita n. sp.* (SI Figure 4). Additionally, we identified, a family of tetraspanins (OG0009162) which are investigated as vaccine candidates in schistosomiasis.(45, 46)

| Category | GO term | p-value | Description |
| --- | --- | --- | --- |
| Cellular component | GO:0005615 | 3.10E-21 | extracellular space |
|  | GO:0000786 | 4.02E-17 | nucleosome |
| Molecular function | GO:0003676 | 1.29E-52 | nucleic acid binding |
|  | GO:0004830 | 1.21E-33 | tryptophan-tRNA ligase activity |
|  | GO:0030527 | 2.78E-18 | structural constituent of chromatin |
|  | GO:0003723 | 6.72E-18 | RNA binding |
| Biological process | GO:0006436 | 1.21E-33 | tryptophanyl-tRNA aminoacylation |
|  | GO:0044571 | 3.17E-19 | [2Fe-2S] cluster assembly |
|  | GO:0051259 | 3.17E-19 | protein complex oligomerization |

“Gene_list_venn_intersection.xlsx”

SI Table 3. Gene list of cross section in Venn diagram. “Gene_list_resistance_associated_literature.xlsx”

SI Table 4. Gene list of resistance associated genes towards albendazole or ivermectin in helminths.

“Orthologous_group_list_duplications.xlsx”

SI Table 5. GO term enrichment analysis results "GO-Term_enrichment_analysis.xlsx"

SI Table 6. Highly duplicated genes. “hits_gwas_all.xlsx”

SI Table 7 “Main_Phenotype_File_sex.txt”

SI Table 8. Worm ID’s with sex for PLINK input and phenotype of drug sensitive or non sensitive: Sex code (’1’ = male, ’2’ = female, ’0’ = unknown), Phenotype value (’1’ = control, ’2’ = case, ’-9’/’0’/non-numeric = missing data if case/control)

